# MACS – a new SPM toolbox for model assessment, comparison and selection

**DOI:** 10.1101/194365

**Authors:** Joram Soch, Carsten Allefeld

## Abstract

**Background:** In cognitive neuroscience, functional magnetic resonance imaging (fMRI) data are widely analyzed using general linear models (GLMs). However, model quality of GLMs for fMRI is rarely assessed, in part due to the lack of formal measures for statistical model inference.

**New Method:** We introduce a new SPM toolbox for model assessment, comparison and selection (MACS) of GLMs applied to fMRI data. MACS includes classical, information-theoretic and Bayesian methods of model assessment previously applied to GLMs for fMRI as well as recent methodological developments of model selection and model averaging in fMRI data analysis.

**Results:** The toolbox – which is freely available from GitHub – directly builds on the Statistical Parametric Mapping (SPM) software package and is easy-to-use, general-purpose, modular, readable and extendable. We validate the toolbox by reproducing model selection and model averaging results from earlier publications.

**Comparison with Existing Methods:** A previous toolbox for model diagnosis in fMRI has been discontinued and other approaches to model comparison between GLMs have not been translated into reusable computational resources in the past.

**Conclusions:** Increased attention on model quality will lead to lower false-positive rates in cognitive neuroscience and increased application of the MACS toolbox will increase the reproducibility of GLM analyses and is likely to increase the replicability of fMRI studies.

## 1 Introduction

In functional magnetic resonance imaging (fMRI), general linear models (GLMs) are routinely applied to relate measured hemodynamic signals with controlled experimental variables in order to make statements about the localization of cognitive functions in brain regions (Friston et al., 1994; Holmes and Friston, 1998). Like any other statistical inference, GLM analyses are dependent on the particular choice of the model (Andrade et al., 1999; Carp, 2012) and model quality can have a profound effect on sensitivity and specificity of statistical tests (Razavi et al., 2003; Monti, 2011).

Model assessment (Akaike, 1974; Schwarz, 1978), model comparison and selection (Kass and Raftery, 1995) as well as model averaging (Hoeting et al., 1999) are established techniques to control for model quality by inferring the most likely model given the data and by conditioning data analysis on the optimal model for each data set. While model assessment (Friston et al., 2007), comparison (Penny et al., 2004; Penny, 2012), selection (Stephan et al., 2009) and averaging (Penny et al., 2010) have become standard practice in dynamic causal modelling (DCM) for fMRI (Friston et al., 2003), such techniques are almost not used in the application of GLMs for fMRI (Friston et al., 1994).

However, formal model inference can be highly beneficial in this context. For example, methodological analyses could concern the inclusion of nuisance regressors such as movement parameters and hemodynamic derivatives (Henson et al., 2002) or physiological recordings (Kasper et al., 2017). Beyond that, empirical analyses could compare prediction error models for reward learning experiments (Abler et al., 2006) or different encoding models for visual reconstruction analyses (Kay et al., 2008).

Importantly, formal model inference does not only enhance the analytical potential of fMRI data analyses, but can also improve their methodological quality. It can be expected that, if models are not selected arbitrarily, but based on objective criteria, this is likely to increase the *replicability* of fMRI studies – i.e. obtaining the same results with the same methods applied to different (or new) data. And, with a validated toolbox for these methods at hand, it will also benefit the *reproducibility* of GLM analyses – i.e. obtaining the same results with the same methods applied to the same data.

Previous attempts to control for model quality in GLMs for fMRI include statistical tests for goodness of fit (Razavi et al., 2003) and the application of Akaike’s or Bayesian information criterion for activation detection (Seghouane and Ong, 2010) or theory selection (Gläscher and O’Doherty, 2010). Additionally, voxel-wise Bayesian model assessment (Penny et al., 2003, 2005, 2007) and random-effects Bayesian model selection (Rosa et al., 2010) have been included in the popular software package *Statistical Parametric Mapping* (SPM), but are only rarely used due to low visibility, high analytical complexity and interpretational difficulty. Finally, a toolbox for frequentist model diagnosis and exploratory data analysis called *SPMd* (“d” for “diagnostic”) has been released for SPM (Luo and Nichols, 2003), but was discontinued several years ago^1^ (Nichols, 2013).

With this work, we provide a new SPM toolbox for **m**odel **a**ssessment, **c**omparison and **s**election termed *MACS* (pronounced as “Max”). Purposes of this software package are to unify classical, information-theoretic and Bayesian measures of model quality previously applied to fMRI data, (ii) to provide a common pipeline for our previously developed methods of cross-validated Bayesian model selection (cvBMS; Soch et al., 2016) and averaging (cvBMA; Soch et al., 2017) that so far existed as seperate toolkits and (iii) to build a framework in which all these model quality measures can be conveniently applied to GLMs estimated in SPM by directly building on `SPM.mat` files.

The rest of the paper falls into three parts. First, we give a technical overview of the features included in the MACS toolbox (see Section 2). Among others, we have implemented goodness-of-fit (GoF) measures, classical information criteria (ICs) and the Bayesian log model evidence (LME). Second, we explain the structure of the MACS toolbox and how it facilitates diverse and user-specific analyses (see Section 3). Most importantly, the embedding of MACS into SPM’s batch systems accelerates and simplies fMRI analyses. Finally, we present two exemplary applications illustrating cvBMS and cvBMA as our latest methodological advances for fMRI (see Section 4) and discuss our work.

## 2 Theory

In the MACS toolbox, methods of model inference are broadly categorized into the three categories *model assessment* (MA), *model comparison* (MC) and *model selection* (MS). This section gives a technical introduction of these different techniques which are also summarized in Table 2.

**Table 1.** Mathematical notation. Note that the number of parameters is equal to the number of GLM regressors *p* in goodness-of-fit measures, but equal to *k* = *p* + 1 in information criteria, because the latter account for the total number of model parameters which include the residual variance *σ*^2^ in addition to the regression coefficients *β*.

| Symbol | general meaning | in GLMs for fMRI |
| --- | --- | --- |
| $m$ | generative model | univariate general linear model (GLM) |
| $\theta$ | model parameters | regression coefficients $\beta$, residual variance $\sigma^2$ |
| $y$ | measured data | blood-oxygenation-level-dependent (BOLD) signal |
| $\hat{\theta}$ | estimated model parameters | estimated regression coefficients $\hat{\beta}$, residual variance $\hat{\sigma}^2$ |
| $\hat{y}$ | predicted data | fitted BOLD signal |
| $n$ | number of data points | number of fMRI scans |
| $p$ | number of model parameters | number of GLM regressors |
| $k$ | | number of GLM regressors, plus one |
| $N$ | number of observations | number of subjects |
| $S$ | | number of sessions |
| $M$ | number of models | number of GLMs |
| $i$ | index over observations | index over scans |
|  |  | index over subjects |
|  |  | index over sessions |
| $j$ | index over models | index over GLMs |
| $y_i$ | a particular observation | the i-th data point |
|  |  | data from the i-th subject |
|  |  | data from the i-th session |
| $m_j$ | a particular model | the j-th GLM |
| $e_j$ | | the j-th GLM (see Fn. 4) |
| $f$ | model family | family of GLMs |

**Table 2.**
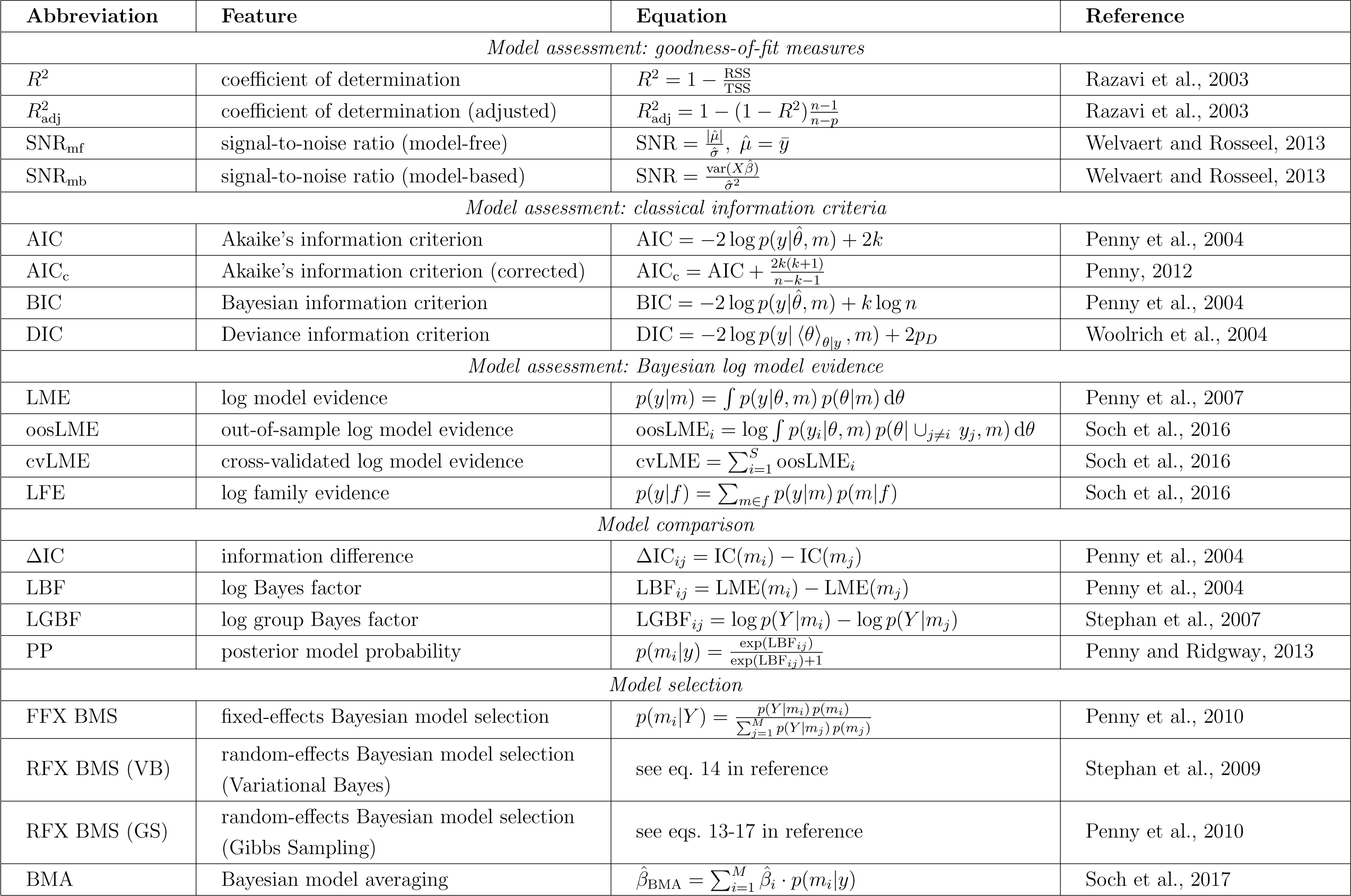
Features in the MACS toolbox. This table summarizes methods of model inference in MACS. References correspond to exemplary applications in the fMRI literature. For the respective statistical literature, confer the list of features (see Section 2).

### 2.1 Model assessment

Model assessment (MA) is defined as any method that quantifies the quality of an individual model by a single real number. MA methods can again be sub-divided into three groups of tools: (i) *goodness-of-fit measures* (GoF) for non-inferential model assessment, (ii) *classical information criteria* (ICs) for frequentist model assessment and (iii) the *log model evidence* (LME) for Bayesian model assessment.

In all of the following, we consider a generative model *m* with model parameters *θ* describing measured data *y* with estimated parameters 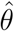 giving predicted data *ŷ*, with number of data points *n* and number of model parameters *p*. The meaning of these variables in the context of GLMs for fMRI is given in Table 1.

#### 2.1.1 Goodness-of-fit measures

The purpose of GoF measures is to get a first impression of how well a model – in this case: a GLM – describes a given set of data – in this case: voxel-wise fMRI signals. One obvious thing to look at would be the *residual variance* of the GLM calculated as

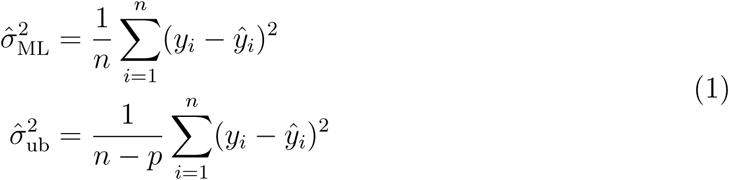

where 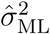 is the maximum-likelihood estimator and 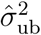 is the unbiased estimator of that quantity. In SPM, the residual variance is calculated somewhat differently as

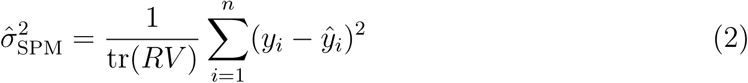

where *R* is the residual forming matrix and *V* is the error correlation matrix of the corresponding GLM which accounts for the loss of effective residual degrees of freedom (erdf) by whitening and filtering the data. Generally, the lower the residual variance, the better model predictions and measured data agree with each other. However, as the residual variance does not have an upper bound, it is more useful to look at the *coefficient of determination* (Razavi et al., 2003, eq. 2)

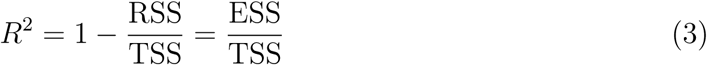

where RSS is the residual sum of squares 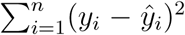, ESS is the explained sum of squares 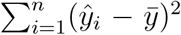 and TSS is the total sum of squares 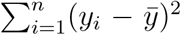, such that TSS = ESS + RSS. Generally, the closer *R*^2^ gets to 1, the better the model captures the variation. However, as the coefficient of determination always increases when another variable is included in the model, an improved measure is given by the *adjusted coefficient of determination* (Razavi et al., 2003, eq. 4)

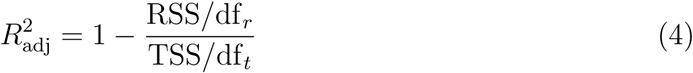

where the residual degress of freedom df_*r*_ = *n* − *p* and the total degrees of freedom df_*t*_ = *n* − 1 are used to adjust the sums of squares to account for additional model parameters. For this reason, the adjusted *R*^2^ can be seen as a precursor to classical information criteria (see next section).

Note that, whereas *R*^2^ always falls between 0 and 1, 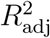 can also become negative. Both *R*^2^ and 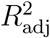 are also related to the *F* -statistic in an *F* -test of a GLM against only the constant regressor which is given by

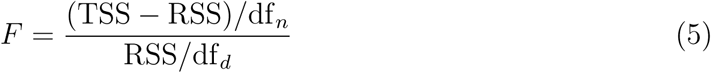

where the numerator degress of freedom df_*n*_ = *p* − 1 and the denominator degrees of freedom df_*d*_ = *n* − *p* are the parameters of the *F* -distribution under the null hypothesis that the signal only consists of a mean *µ*.

Goodness-of-fit measures also include the *signal-to-noise ratio* (SNR):

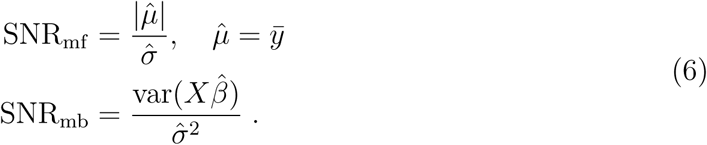

Here, SNR_mf_ is the model-free SNR which is calculated as the inverse of the *coefficient of variation* (CoV) and quantifies the ratio of signal average to signal variation (Welvaert and Rosseel, 2013, def. 1). Signals that are very low or very variable will receive a lower SNR given this measure. In contrast, SNR_mb_ corresponds to a model-based SNR which divides the variance of the explained signal by the variance of the unexplained signal (Welvaert and Rosseel, 2013, def. 5).

Note that, while both measures are unit-less, they cannot be directly compared. Generally, with a change of the GLM, the model-based SNR will also change while the modelfree SNR will not. Moreover, for a mean-only GLM, the model-based SNR will be zero (because a constant regressor has no variation) while the model-free SNR will be non-zero (given that the BOLD signal has a non-zero mean).

Also observe that, if the measured signal *y* and the predicted signal *ŷ* have the same mean, then 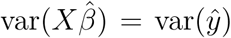 is proportional to the explained sum of squares and the SNR simplifies to SNR_mb_ = ESS*/*RSS = *R*^2^*/*(1 − *R*^2^).

In our toolbox, all these measures are implemented via the routine `MA_inspect_GoF` and can be exported as whole-brain maps or explored in a voxel-wise manner (see Figure 4A).

#### 2.1.2 Classical information criteria

Classical information criteria (ICs), in the terminology of the MACS toolbox, are model quality measures that only depend on the *maximum log-likelihood* (MLL), as a measure of model accuracy, as well as *n* and *p*, giving rise to some measure of model complexity. The first of these ICs was *Akaike’s information criterion* (Akaike, 1974)

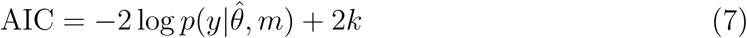

where the maximum log-likelihood 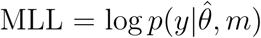 and *k* = *p* + 1 is the number of model parameters (see Table 1). The *corrected AIC*, an improved version of the AIC for finite-sample calculations, has been derived as (Hurvich and Tsai, 1989)

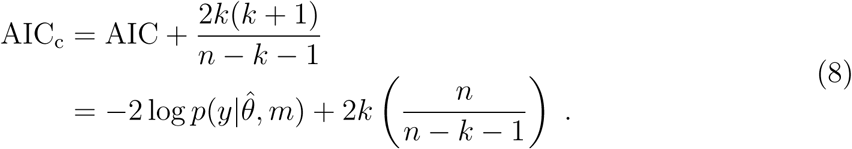

Because the corrected AIC improves the AIC for small samples and approximates the AIC for large samples with *n* ≫ *k*, it can in principle always be used in place of the AIC. Historically, the second IC was the *Bayesian information criterion* (Schwarz, 1978)

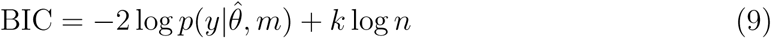

which means that AIC and BIC are equal for sample sizes around *n* = *e*^2^ ≈ 7.39. Although the name suggests that the BIC rests on Bayesian inference, it does not and is only theoretically connected to Bayesian statistics by being an infinite-sample approximation to the Bayesian log model evidence (see next section).

In contrast to that, the *deviance information criterion* (DIC) is a “real” Bayesian information criterion, because it requires the specification of prior distributions on the model parameters and posterior distributions are used to calculate expected likelihoods. Technically, the DIC therefore depends on more than just MLL, *n* and *p*. Formally, the DIC is given by (Spiegelhalter et al., 2002)

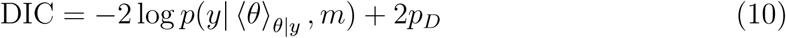

where the effective number of parameters *p*_*D*_ is

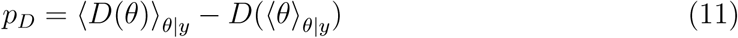

and *D*(*θ*) is the log-likelihood function on the deviance scale^2^

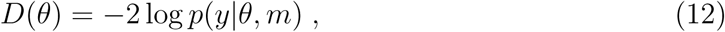

such that ⟨*D*(*θ*)⟩_*θ*|*y*_ is proportional to the *posterior expected log-likelihood* (PLL) and *D*(⟨*θ* ⟩_*θ*|*y*_) is proportional to the *log-likelihood at the posterior expectation* (LLP). In these equations, the operator ⟨ ·⟩_*θ*|*y*_ denotes an expectation across the posterior distribution (Woolrich et al., 2004) which naturally depends on the prior distribution. However, the DIC is very robust with respect to the choice of the prior and we have therefore implemented it with a non-informative prior (Soch et al., 2016, eq. 15).

In our toolbox, all these criteria are implemented via the routine `MA_classic_ICs` which can also be accessed through the corresponding batch editor job (see Figure 2).

#### 2.1.3 Bayesian log model evidence

An alternative to those classical information criteria are Bayesian measures of model quality based on the marginal log-likelihood, also called the *log model evidence* (LME) which is given by (Gelman et al., 2013, eq. 1.3; Bishop, 2007, eq. 1.45)

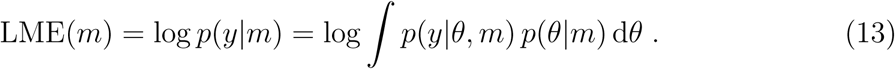

Note that, in contrast to GoF measures and classical ICs which use point estimates for model parameters, the LME calculates an expectation over the entire parameter distribution. Furthermore, while classical ICs operate on the deviance scale (see eq. 12), the LME operates on the log-likelihood scale (see eq. 13), so that thresholds commonly applied to LME differences (Kass and Raftery, 1995), i.e. log Bayes factors (see next section), must be multiplied by −2 when applying them to IC differences.

The LME has several advantages over classical information criteria including (i) its automatic penalty for additional model parameters (Penny, 2012), (ii) its natural decomposition into model accuracy and model complexity (Penny et al., 2007) and (iii) its consideration of the whole uncertainty about parameter estimates (Friston et al., 2007). The LME however has the disadvantage that it depends on a prior distribution *p*(*θ*|*m*) over model parameters and the choice of the prior can have a profound impact on resulting model comparisons. In recent work on GLMs for fMRI, we have therefore suggested the *out-of-sample log model evidence* (Soch et al., 2016, eq. 13)

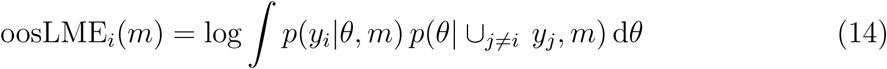

and the *cross-validated log model evidence* (Soch et al., 2016, eq. 14)

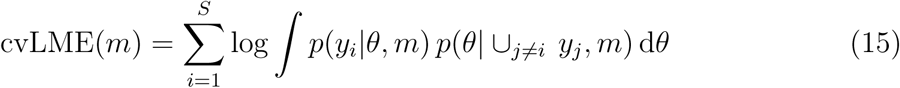

where *S* is the number of data subsets – in our case: fMRI recording sessions^3^. These can be seen as Bayesian measures of model quality, but avoid having to specify prior distributions on the model parameters. Instead, the prior distribution is obtained by training the model on independent data of the same kind.

Given that LME, cvLME or oosLME have been calculated and models *m* can be sensibly grouped into model families *f, log family evidences* (LFE) can be easily calculated using the law of marginal probability as (Soch et al., 2016, eq. 18)

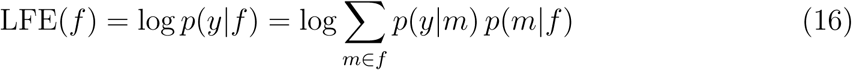

where *p*(*y*|*m*) is the exponential of LME, cvLME or oosLME and *p*(*m*|*f*) is a within-family prior, usually set to a discrete uniform distribution, embodying the assumption that within each family, all the models are equally likely *a priori*.

In our toolbox, the cvLME is the default model selection criterion and can be calculated via the routine `MA_cvLME_multi` or the corresponding batch editor job (see Figure 2).

### 2.2 Model comparison

Model comparison (MC) is defined as any method that quantifies the quality difference between two models by a single real number. Most trivially, one can for example calculate the *difference between two information criteria* (Penny et al., 2004, eq. 23)

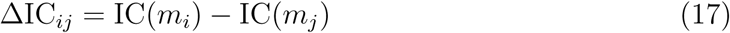

which can be performed for AIC, BIC etc. and compared with particular scales. When calculating this difference between two LMEs, the result is called the logarithm of a Bayes factor or a *log Bayes factor* (Penny et al., 2004, eq. 22)

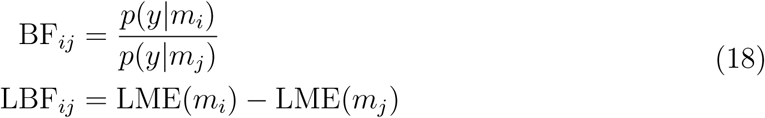

which can be applied to LME, cvLME or oosLME. There are two well-known scales of interpretation (Jeffreys, 1961; Kass and Raftery, 1995) which e.g. define thresholds for LBFs of 0, 1, 3 or 5 corresponding to BFs of around 1, 3, 20 or 150 and evidence in favor of model *m*_*i*_ against model *m*_*j*_ labeled as “not worth more than a bare mention”, “positive”, “strong” or “very strong” (Kass and Raftery, 1995, p. 777). While these scales belong to standard practice in Bayesian inference, we strongly advocate to always also report concrete LBF values and not to misunderstand LBFs exceeding a threshold as signalling “statistical significance” (McShane et al., 2017).

When analyzing a group of subjects in an experimental study and assuming independence across subjects, subject-level log model evidences can be summed up into a *group-level log model evidences* (Penny et al., 2010, eq. 6)

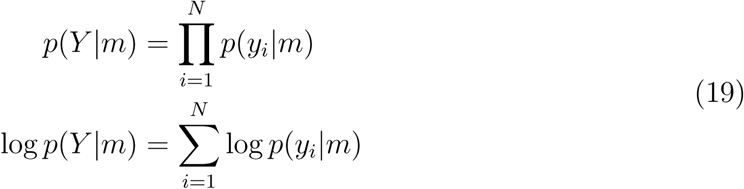

where *Y* = {*y*_1_, *…, y*_*N*_} are the group data and *N* is the number of subjects. Comparing two models across several subjects then gives rise to a group Bayes factor or *log group Bayes factor* which is given by (Stephan et al., 2007, eq. 20)

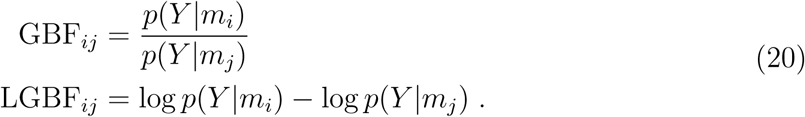

Importantly, this LGBF approach assumes the same model for each subject and therefore corresponds to a fixed-effects (FFX) analysis. If this assumption cannot be justified, a random-effects (RFX) model has to be used (see next section).

Finally, in the case that just two models are considered and a uniform prior across models is assumed, (log) Bayes factors can be directly transformed into *posterior model probabilities* using the simple relation (Penny and Ridgway, 2013, eq. 10)

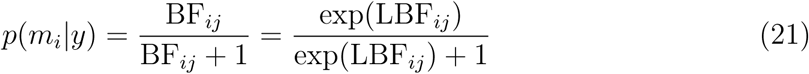

and additionally

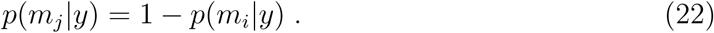

### 2.3 Model selection

Model selection (MS) is defined as any method that chooses one particular model from a set of candidate models based on some objective criterion. For example, *fixed-effects Bayesian model selection* (FFX BMS) generalizes the idea of posterior model probabilities to more than two models, such that (Penny et al., 2010, eq. 7)

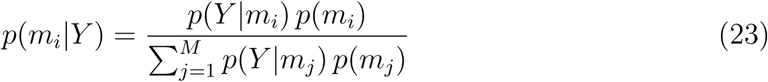

where *M* is the number of models. The optimal model is then the one with the highest posterior probability. Note that, by using the group-level LME *p*(*Y*|*m*_*i*_), FFX BMS effectively assumes the same optimal model for all subjects.

In contrast to that, *random-effects Bayesian model selection* (RFX BMS) allows for the possibility that different subjects have different optimal models. For each subject’s data *y*_*i*_, the generating model *m*_*i*_ is assumed to come from a multinomial distribution with a Dirichlet as the prior distribution over multinomial frequencies *r*. However, the models are not observed directly, but only indirectly via the model evidences from first-level model assessment^4^ (Stephan et al., 2009, eqs. 3-5):

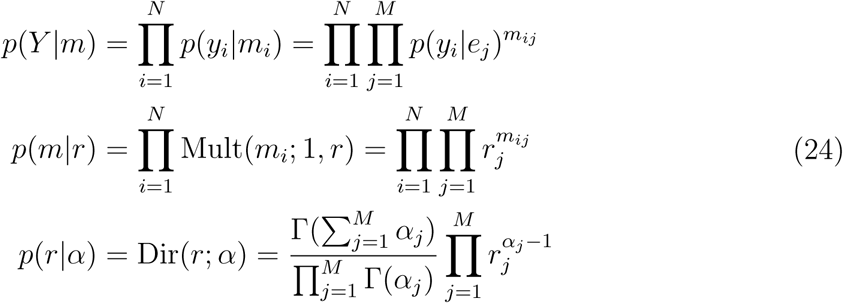

Together, these probability distributions define a second-level hierarchical population proportion model that can be inverted using either Variational Bayes (VB) (Stephan et al., 2009, eq. 14) or Gibbs Sampling (GS) (Penny et al., 2010, eqs. 13-17) techniques. When using model selection for methodological control, i.e. identifying the optimal model for data analysis, the optimal model is simply the one with the highest estimated model frequency, quantified via e.g. expected frequencies or likeliest frequencies (Soch et al., 2016, p. 474). In contrast, when the purpose is discovery science, i.e. deciding between competing theories of brain function, the best model should outperform the others considerably, quantified using e.g. exceedance probabilities (Stephan et al., 2009, eq. 16).

Finally, *Bayesian model averaging* (BMA) is also grouped under model selection here as it implicitly consists in, based on the set of candidate models, fitting a larger model which effectively marginalizes over modelling approaches. In our implementation, averaged model parameter estimates are calculated as (Soch et al., 2017, eq. 8)

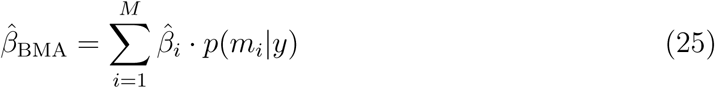

where 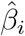 is the *i*-th model’s parameter estimate for a particular regressor in the GLM and *p*(*m*_*i*_|*y*) is the *i*-th model’s posterior probability calculated from cvLMEs.

Whereas *model selection* decides for one particular model that maximizes a certain function such as log model evidence, posterior model probability or group-level model frequency, *model averaging* combines several models using their posterior probabilities calculated from model evidences. Given that finite data never provide absolute certainty about the modelling approach, this approach allows to account for modelling uncertainty by utilizing parameter estimates from all models in the model space instead of discarding the valuable information from all but one model (Soch et al., 2017, p. 188).

## 3 Methods

The MACS toolbox is written in MATLAB and designed to work as an extension for Statistical Parametric Mapping (SPM), Version 8 or 12 (SPM8/12). This section describes how three layers of implementation make the toolbox accessible for different users and how two modes of batch processing make the toolbox useful in different contexts.

### 3.1 Three layers of implementation

An overview of the toolbox functions is given in Figure 1. This figure shows the three types of functions included in the toolbox: (i) mathematical functions, (ii) interface functions and (iii) batch functions. *Mathematical functions* (see Figure 1C) directly implement the statistical analyses (such as `ME_GLM_NG` for estimating a Bayesian GLM) required for toolbox features. *Interface functions* (see Figure 1B) represent toolbox features (such as `MA_cvLME_multi` for calculating the voxel-wise cvLME) and execute mathematical functions. *Batch functions* (see Figure 1A) take information from SPM batch editor jobs (such as `batch_MA_cvLME_auto` for performing cvLME assessment) and use it to call interface functions.

**Figure 1.**
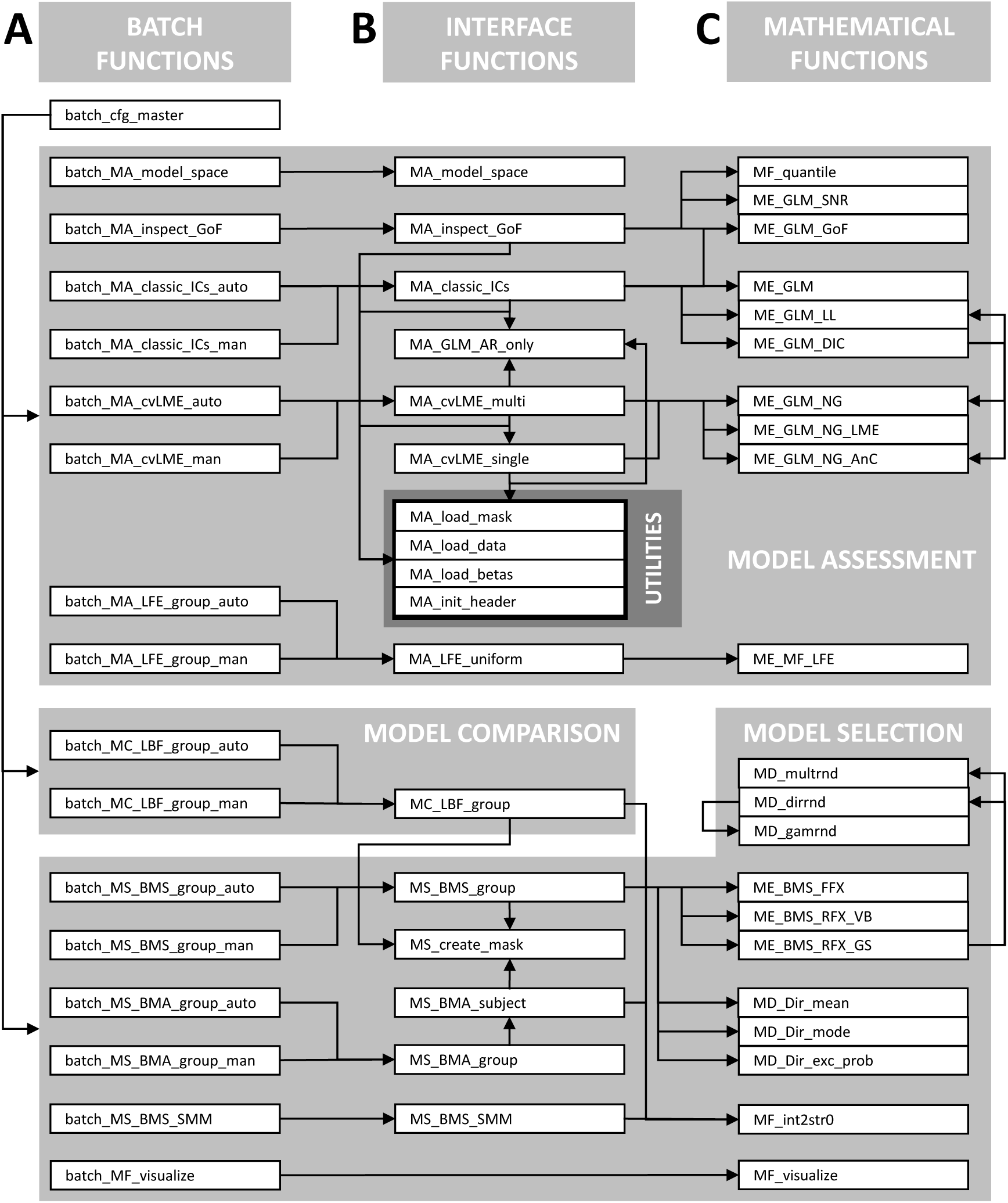
Architecture of the MACS toolbox. This figure is for developers only and lists all functions from the toolbox. A black arrow indicates that one function calls the other. The horizontal dimension corresponds to the type of function (see Section 3.1): Generally, (A) batch functions call (B) interface functions which in turn call (C) mathematical functions. The vertical dimension corresponds to the type of feature (see Section 2): Interface functions consist of tools for model assessment (function prefix “MA”), model comparison (MC) and model selection (MS); mathematical functions consist of tools for model estimation (ME), many distributions (MD) and more functions (MF).

**Figure 2.**
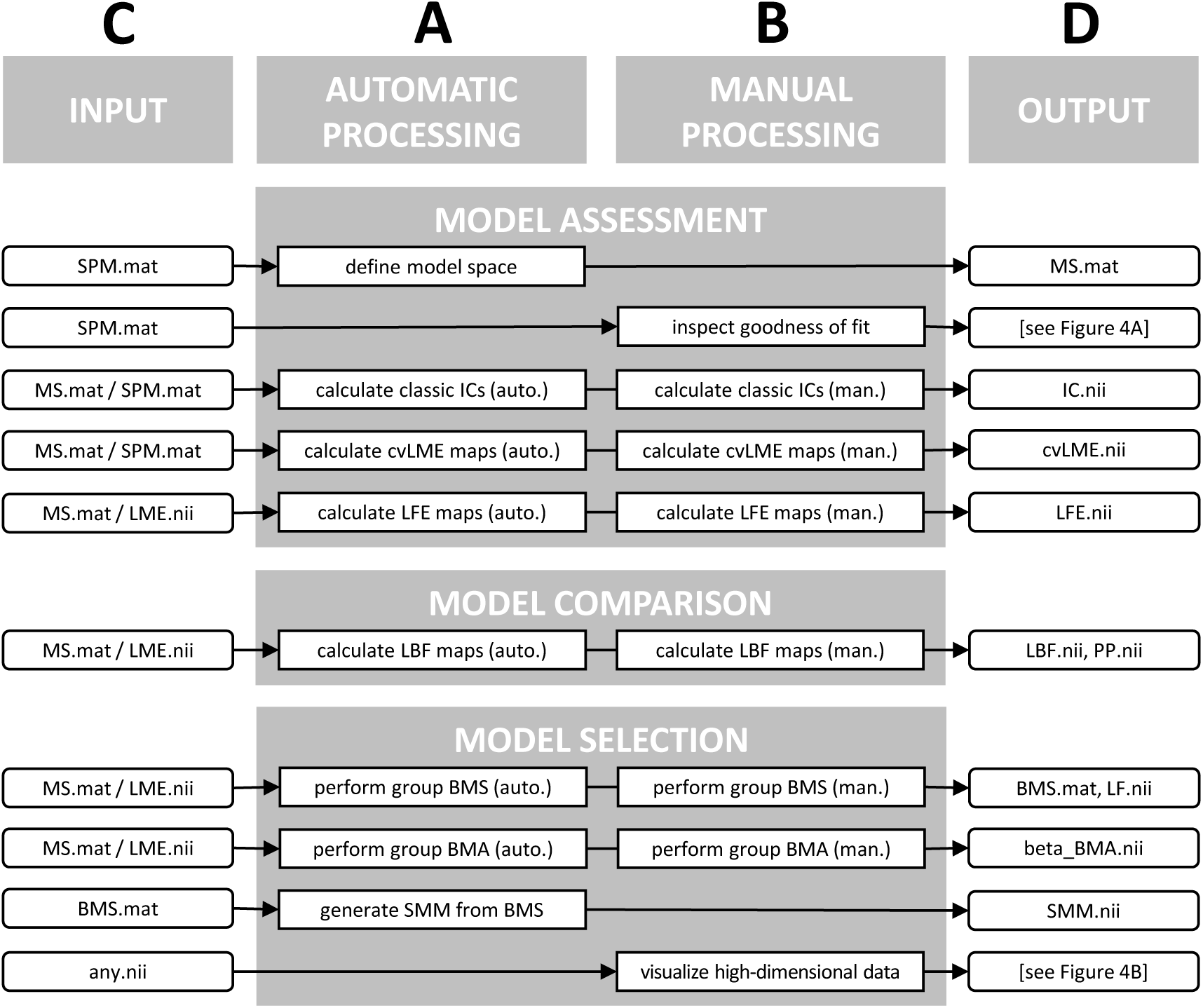
Batch modules in the MACS toolbox. This figure lists batch modules from the toolbox that are accessible via the SPM batch editor. Modules either belong to (A) the automatic processing stream or (B) the manual processing stream (see Section 3.2): automatic processing is based on a model space file (`MS.mat`) whereas manual processing is based on manual input of GLM info (`SPM.mat`) or cvLME maps (`LME.nii`). Modules take (C) particular input files and result in (D) particular output files: MATLAB files (`.mat`) contain structured information (such as GLM info) and NIfTI files (`.nii`) are brain images (such as cvLME maps). For abbreviations in filenames, confer the list of features (see Section 2 and Table 2).

*Regular toolbox users* will never access any of these functions directly in MATLAB, but will use toolbox features through SPM’s batch editor only. This can be achieved by installing the MACS toolbox and opening the SPM batch editor (see Section 6 and Figure 3). Users can then fill out regular batch jobs like in SPM and directly run them from the batch editor. Batch job templates only use few options most of which can often be left at their default values. Examples are given in Figures 3 and 4:

**Figure 3.**
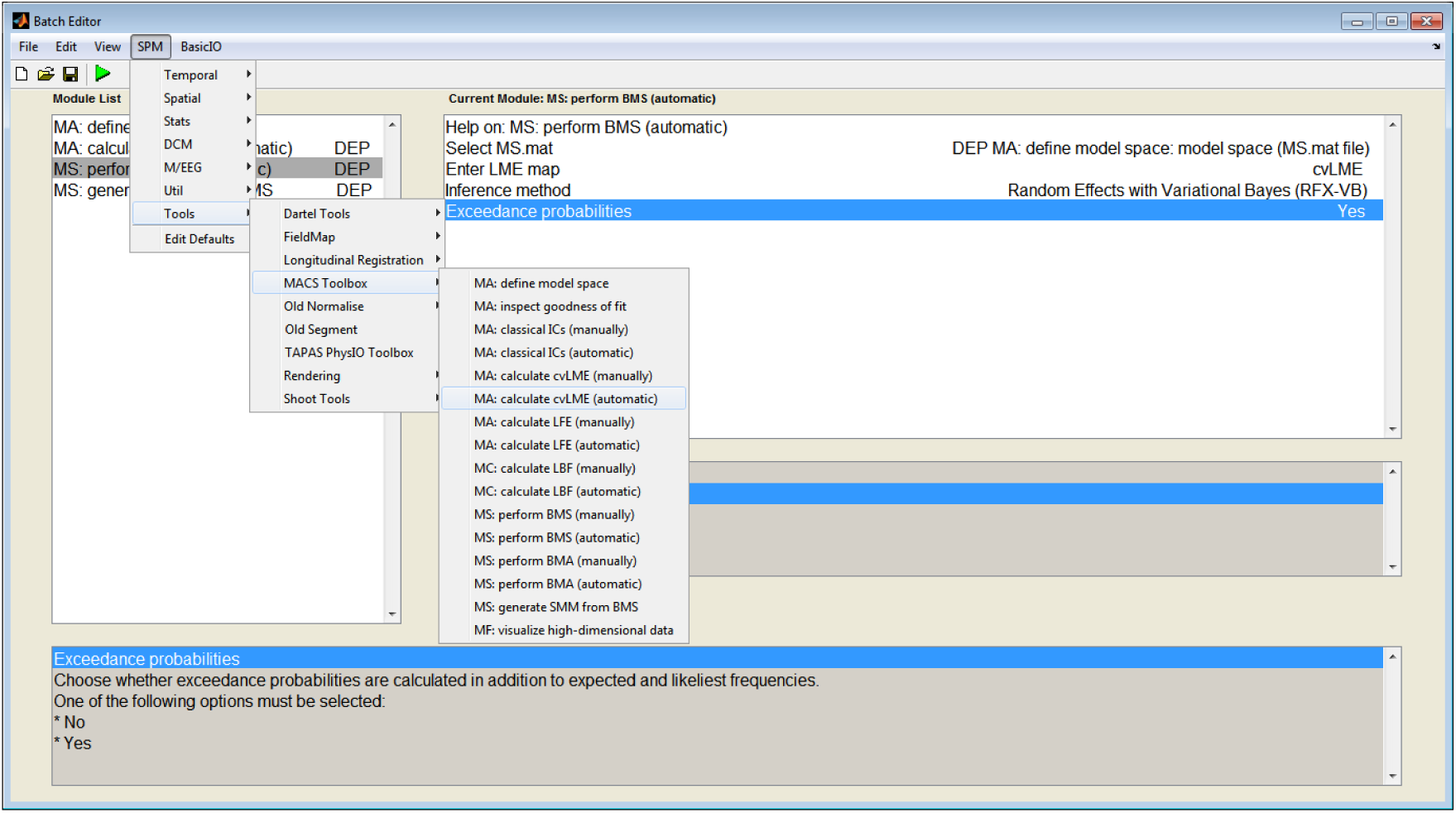
Exemplary batch for Bayesian model selection. This figure shows the SPM batch editor with a typical batch for cross-validated Bayesian model selection (cvBMS) including an active module for group-level Bayesian model selection (BMS). The unfolded menu also hints at where to find features from the MACS toolbox upon successful integration into the SPM software, with the module for automatic cvLME model assessment being highlighted by mouse-over.

~~~
- Figure 3 : batch for cross-validated Bayesian model selection
- Figure 4A: output from goodness-of-fit inspection for a GLM
- Figure 4B: output from high-dimensional data visualization
~~~

*Intermediate toolbox users* may also use the batch editor, but will additionally call the interface functions via MATLAB’s command-line interface. This could for example happen in a pipeline script which is intended to process many models applied to many subjects. Note however that those pipeline-style analyses can also be implemented using the regular batch system. An exemplary command-line call could look like this:

~~~
>> load […]\SPM.mat        % load GLM estimated in SPM
>> MA_cvLME_multi(SPM)     % calculate voxel-wise cvLME
>> MA_classic_ICs(SPM)     % calculate voxel-wise AIC/BIC
~~~

**Figure 4.**
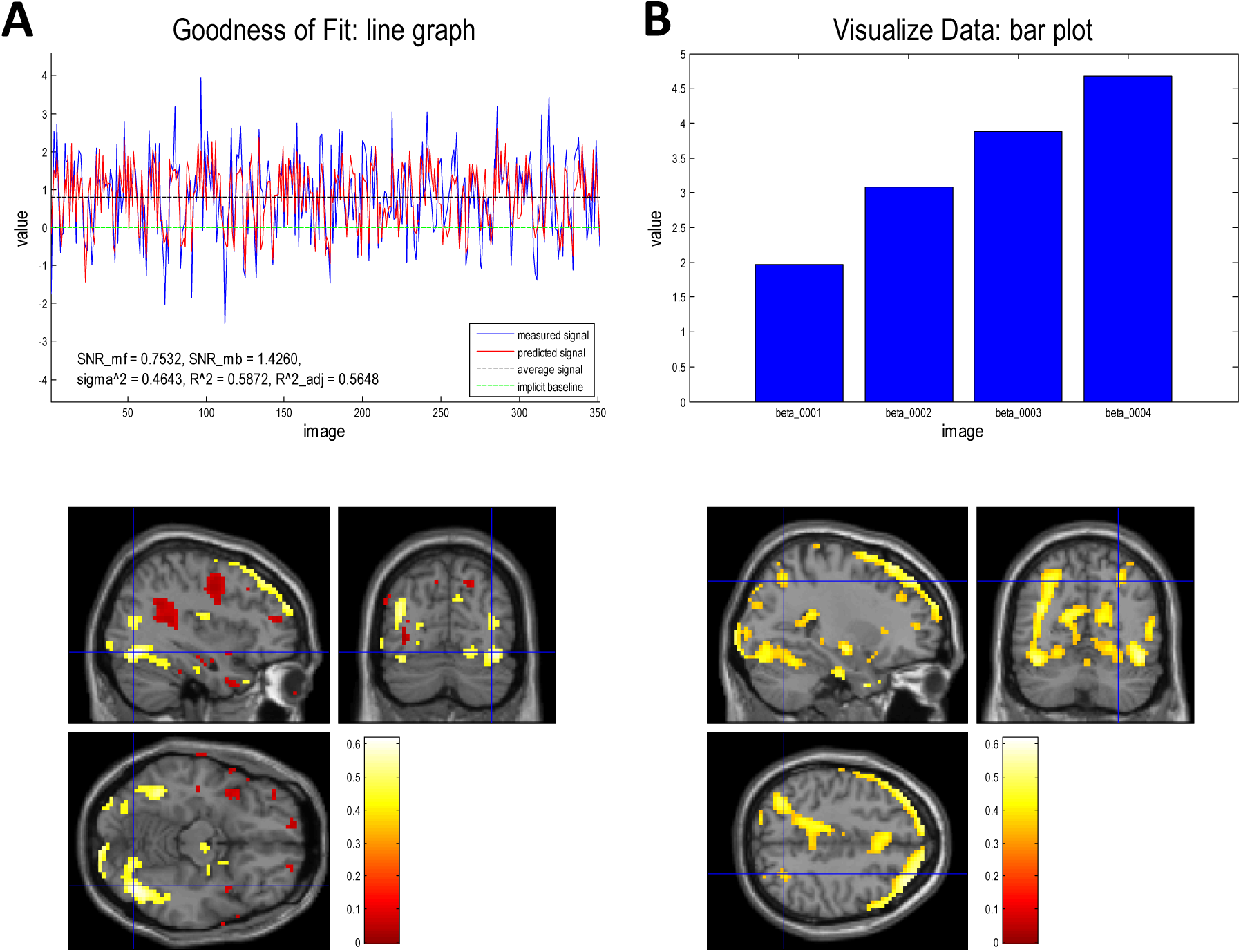
Demonstrations of voxel-wise visualization. This figure shows SPM graphics windows resulting from voxel-wise visualization tools included in the MACS toolbox. (A) Using the module “inspect goodness of fit”, measured and predicted fMRI signal are shown alongside goodness-of-fit measures (see Section 2.1.1) which can be browsed for every in-mask voxel included in a particular first-level analysis (see Ashburner et al., 2016, sec. 31.2). The lower panel highlights voxels with the top 5% (yellow, good fit) and the bottom 5% (red, bad fit) in terms of coefficient of determination (*R*^2^) for this GLM. (B) Using the module “visualize high-dimensional data”, several data points in each brain region can be displayed and browsed in a voxel-wise fashion. The lower panel highlights voxels in which the *R*^2^ of a GLM (the same as in A) is larger than 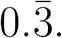. The upper panel bar plots beta estimates from the four experimental condition regressors of this model, indicating a negative effect of both two-level factors in this voxel (see Ashburner et al., 2016, fig. 31.1). Note that this feature may also be used to display voxel-wise posterior probabilities or model frequencies in large model spaces for comparing model preferences in different regions of interest.

*Advanced toolbox users* and developers can also rely on the batch editor and interface functions, but will additionally call mathematical functions to perform their own customized analyses. This could for example happen when one is interested in Bayesian GLM parameter estimates or when the cvLME is to be calculated for a large number of models in a small number of voxels. This type of usage can also lead to new methods developments. An extract from a possible script could read like this:

~~~
01 % load model specification and prepare model estimation
02 load […]\SPM.mat           % load GLM estimated in SPM
03 Y = MA_load_data(SPM);     % load voxel-wise fMRI data
04 X = SPM.xX.X;              % get design matrix
05 V = SPM.xVi.V;             % get covariance matrix
06 P = spm_inv(V);            % get precision matrix
07 [n, p] = size(X);          % get model dimensions
08 m0 = zeros(p,1);           % set prior beta mean
09 L0 = zeros(p,p);           % set prior beta covariance
10
11 % estimate general linear model with normal-gamma priors
12 [mn, Ln, an, bn] = ME_GLM_NG(Y, X, P, m0, L0, 0, 0);
13 LME = ME_GLM_NG_LME(P, L0, 0, 0, Ln, an, bn);
~~~

### 3.2 Two modes of batch processing

An overview of the toolbox modules is given in Figure 2. This figure shows the two modes of batch processing that the toolbox allows: (i) automatic processing and (ii) manual processing. In short, automatic processing requires specifying the model space only once whereas manual processing requires that log model evidence images are entered for each model and each subject in every processing step.

*Automatic processing* consists in using the toolbox module for model space definition to create a model space file called `MS.mat` that contains references to `SPM.mat` files belonging to several GLMs for several subjects. Given that the model space has been specified, cvLME model assessment, Bayesian model selection and other modules can be implemented by a simple reference to the model space file which automatically applies the analyses to all subjects. The automatic procesing mode is the prefered one as it allows larger analyses in shorter time and also avoids unnecessary analysis errors.

*Manual processing* means that GLM folders or cvLME maps are not referenced automatically using a model space file, but manually by reference to their filepaths. This is useful when for example cvLME model assessment is prefered to be performed in one batch with GLM model estimation or when LME images are processed, e.g. spatially smoothed or recalculated into LFE images, before further analysis steps are performed.

Consider the example of cross-validated Bayesian model selection (cvBMS; Soch et al., 2016): In the automatic processing mode, each model for each subject would have to be referenced *once* – using its `SPM.mat` when defining the model space. In the manual processing mode, each model for each subject would have to be referenced *twice* – using its `SPM.mat` when calculating the cross-validated log model evidence and using the resulting `LME.nii` when performing second-level model selection.

Like the SPM batch system, the MACS toolbox uses *dependencies between modules*. Dependencies are references to files generated in earlier processing steps that are to be used in later processing steps. For example, an `SPM.mat` file generated during model estimation may be used for cvLME calculation; or a cvLME map generated during model assessment may be used for group-level BMS; or a `BMS.mat` file generated during model selection may be used for generation of selected-model maps (SMM); etc. The MACS toolbox encourages the use of dependencies and they are most conveniently implemented in the automatic processing mode.

## 4 Results

The MACS toolbox was primarily designed to summarize tools for *cross-validated Bayesian model selection* (cvBMS) and *cross-validated Bayesian model averaging* (cvBMA). In this section, we propose pipelines and analyze examples to demonstrate how these techniques can be effortlessly implemented with MACS. Before that, we describe analyses illustrating how to apply goodness-of-fit measures and classical information criteria to first-level GLMs in SPM.

### 4.1 Model assessment: repetition priming

To demonstrate first-level model assessment, we reanalyze a single-subject, single-session data set from a study on repetition priming (Henson et al., 2002) that is used as example data in the SPM manual (Ashburner et al., 2016, ch. 31). In this experiment, subjects were viewing faces which either belonged to famous or non-famous people (“familiarity”) and each face was presented twice (“repetition”). This implies a 2 × 2 experimental design with 4 experimental conditions. This demonstration builds on data preprocessed according to the SPM12 manual and addresses model assessment for the “categorical” model and the “parametric” model (Ashburner et al., 2016, sec. 31.1-3).

First, we want to get an impression of where in the brain the categorical model performs better and where it performs worse. We use the MACS module for goodness-of-fit inspection to assess signal-to-noise ratio (SNR) and coefficient of determination (*R*^2^) across the whole brain. We find that in visual cortex, especially fusiform face area (FFA), the model achieves the highest variance explanation which is consistent with the facial stimuli of the experiment (see Figure 4A).

Second, we want to find out where the model fit is supported by which model parameters. We use the MACS module for high-dimensional data visualization to explore the estimated regression coefficients from the main experimental conditions in a voxel-wise fashion. For orientation purposes, we overlay this query with an *R*^2^ image obtained in the first step. We find that there is one region of parietal cortex, possibly the superior parietal gyrus (SPG), in which there is an effect of both experimental factors, familiarity and repetition (see Figure 4B).

Finally, we want to compare the categorical against the parametric GLM. We use the MACS module for classical information criteria (ICs) to compute the voxel-wise Bayesian information criterion (BIC) for both models. Computing the information difference (ΔIC) between the two models is then implemented using the log Bayes factor (LBF) module. Model comparison at an ad-hoc threshold of ΔIC < −10 in favor of the parametric model^5^ gives rise to a large cluster in parietal cortex, close to the postcentral gyrus (PCG), that was also reported when performing an *F* -test on the additional regressors in the parametric model^6^ (Ashburner et al., 2016, fig. 31.17).

### 4.2 Model selection: orientation pop-out

To demonstrate the cvBMS pipeline, we reanalyze data from an earlier study (Bogler et al., 2013) that has previously been used for the cvBMS paper (Soch et al., 2016). In this data set, we compared three models: a categorical model (GLM I) that describes the 4 × 4 experimental design using 16 regressors, one for each experimental condition; a linear-parametric model (GLM II) that uses one onset regressor indicating stimulation phases (orientation pop-out) and two parametric modulators encoding levels of the two factors (left and right orientation contrast); and a nonlinear-parametric model (GLM III) that uses nonlinear parametric regressors, motivated by behavioral pre-tests (Bogler et al., 2013; Soch et al., 2016). In this demonstration, we will build on estimated first-level GLMs and only describe the steps of model assessment and model selection.

First, in order to avoid model dilution, we want to compare the family of categorical models (consisting of GLM I only) against the family of parametric models (consisting of GLMs II & III). To this end, we specify a job in which we define a model space that consists of all 3 models applied to 22 subjects. Using dependencies on this model space information, automatic calculation of cvLME maps and LFE maps is performed (see Figure 5A). Following model family assessment, group-level BMS has to be specified manually, because log *family* evidences are not associated to particular *models* – which would be necessary for the automatic processing mode –, but only exist as individual images. After group-level BMS, selected-model maps are generated (see Figure 5A). This analysis leads to the results depicted in Figure 3B of the cvBMS paper.

**Figure 5.**
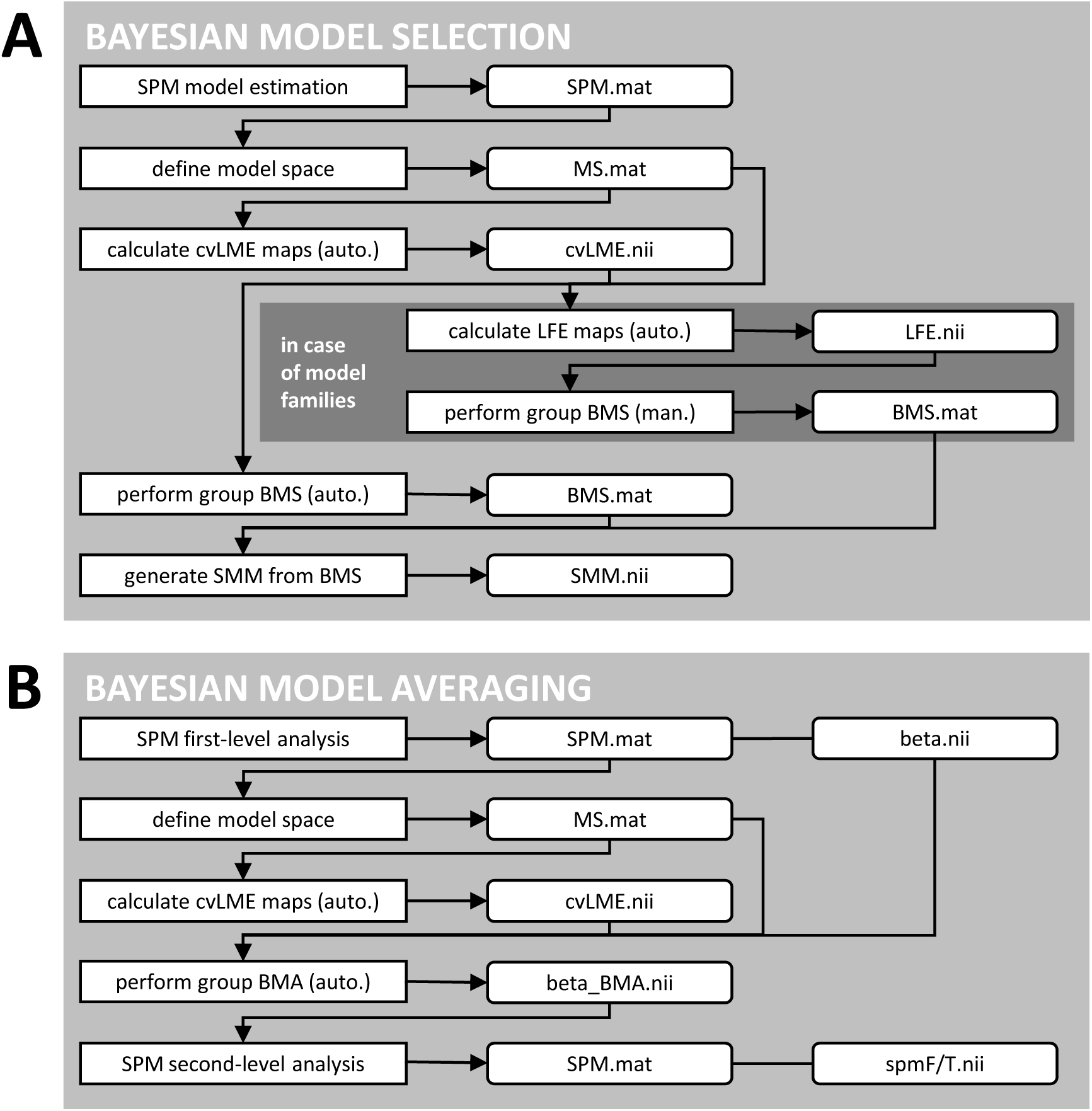
Suggested pipelines for the MACS toolbox. This figure shows processing pipelines underlying the examples reported in this paper (see Section 4). (A) For Bayesian model selection (BMS), SPM model estimation should be followed by definition of the model space, automatic calculation of cvLME maps, automatic group-level BMS and generation of selected-model maps (SMM) from BMS results. In the case that individual models can be grouped into model families, automatic calculation of LFE maps would be followed by manual group-level BMS. (B) For Bayesian model averaging (BMA), SPM first-level analysis should be followed by definition of the model space, automatic calculation of cvLME maps, automatic group-level BMA which invokes subject-level BMAs (see Figure 1) and SPM second-level analysis based on averaged parameters.

Second, within the winning family of parametric models, we want to compare the linear-parametric (GLM II) to the nonlinear-parametric (GLM III) model. To this end, we define a new model space consisting only of these 2 models applied to 22 subjects. As cvLME maps have already been generated in the first step, model assessment does not have to be performed in this second step. Using dependencies on model space information, automatic group-level BMS and generation of SMMs are performed (see Figure 5A). This analysis leads to the results depicted in Figure 3C of the cvBMS paper.

Third, using the winning nonlinear-parametric model, we want to investigate the effect of removing the parametric modulator either for left or for right orientation contrast (OC), i.e. the effect of visual hemifield on model preference. This leads to a model describing only left OC (GLM III-l) and a model describing only right OC (GLM III-r). Again, we define a model space consisting of these 2 models applied to 22 subjects. As these are new models, cvLME maps have to be calculated (automatically). Finally, group-level BMS and selected-model maps are added to the pipeline (see Figure 5A). From the SMMs, we extract the proportion of voxels in left and right V4, respectively, with preferences for left or right OC, respectively, according to the group-level model selection. This analysis leads to the results depicted in Figure 5C of the cvBMS paper.

### 4.3 Model averaging: conflict adaptation

To demonstrate the cvBMA pipeline, we reanalyze data from another study (Meyer and Haynes, in prep.) that has previously been used for the cvBMA paper (Soch et al., 2017) which also details the experimental paradigm, stimulus material and data preprocessing. For this data set, we estimated four models: all models include regressors for main blocks and delay phases in a conflict adaptation paradigm (Meyer and Haynes, in prep.); one model includes no additional processes (GLM 1); one model additionally describes the first trials following preparatory delays (GLM 2); one model additionally describes the cue phases preceding preparatory delays (GLM 3); and one model includes both additional processes, cue phases and first trials (GLM 4). Critically, cue phases and first trials are temporally correlated to preparatory delays, such that model choice influences parameter estimates for the delays phases. However, as each model includes regressors for the delay phases, we can use Bayesian model averaging to remove this modelling uncertainty (Soch et al., 2017). In this demonstration, we will build on estimated first-level GLMs and only describe the steps of model assessment and model averaging.

First, following model estimation, we want to calculate averaged estimates for the common model parameters in these four GLMs. To this end, we specify a job in which we define a model space that consists of all 4 models applied to 24 subjects. Using dependencies on this model space information, automatic calculation of cvLME maps and automatic group-level BMA are performed (see Figure 5B). Additionally, we perform automatic group-level BMS and generate selected-model maps. This analysis leads to the results reported in Table 2 and Figure 4A of the cvBMA paper. Group-level SMMs can be used for masking second-level analyses – an alternative to BMA that is not included in this demonstration, but discussed in the publication (Soch et al., 2017).

Second, following model averaging, we want to perform second-level analysis, but based on the averaged model parameter estimates calculated in the first step instead of the parameter estimates from a particular GLM as in standard processing pipelines. To this end, we use SPM’s factorial design specification to set up a paired t-test between parameter estimates belonging to the two delay phase regressors (see Figure 5B). Subsequent model estimation, contrast calculation and statistical inference lead to the results reported in Table 1 and Figure 4A of the cvBMA paper. As expected, a main effect between switch and stay delays phases and, in more detail, a positive effect of switch over stay delay phases can be observed in left primary motor cortex (Soch et al., 2017).

In addition to a PDF of the user manual for the MACS toolbox, SPM batch editor job files from all demonstrations, classical model assessment as well as cvBMS and cvBMA, are included as MAT files in the MACS toolbox repository. They can be used for educational purposes illustrating suggested pipelines or as analysis templates underlying the user’s applications. Additionally, time consumption of creating and executing these demonstrations are reported in Table 3. In summary, when using the automatic processing mode, complex analyses, such as whole cvBMS and cvBMA pipelines, can be quickly implemented, because almost all analysis steps can simply reference the model space module which usually is the only time-consuming operation.

**Table 3.**
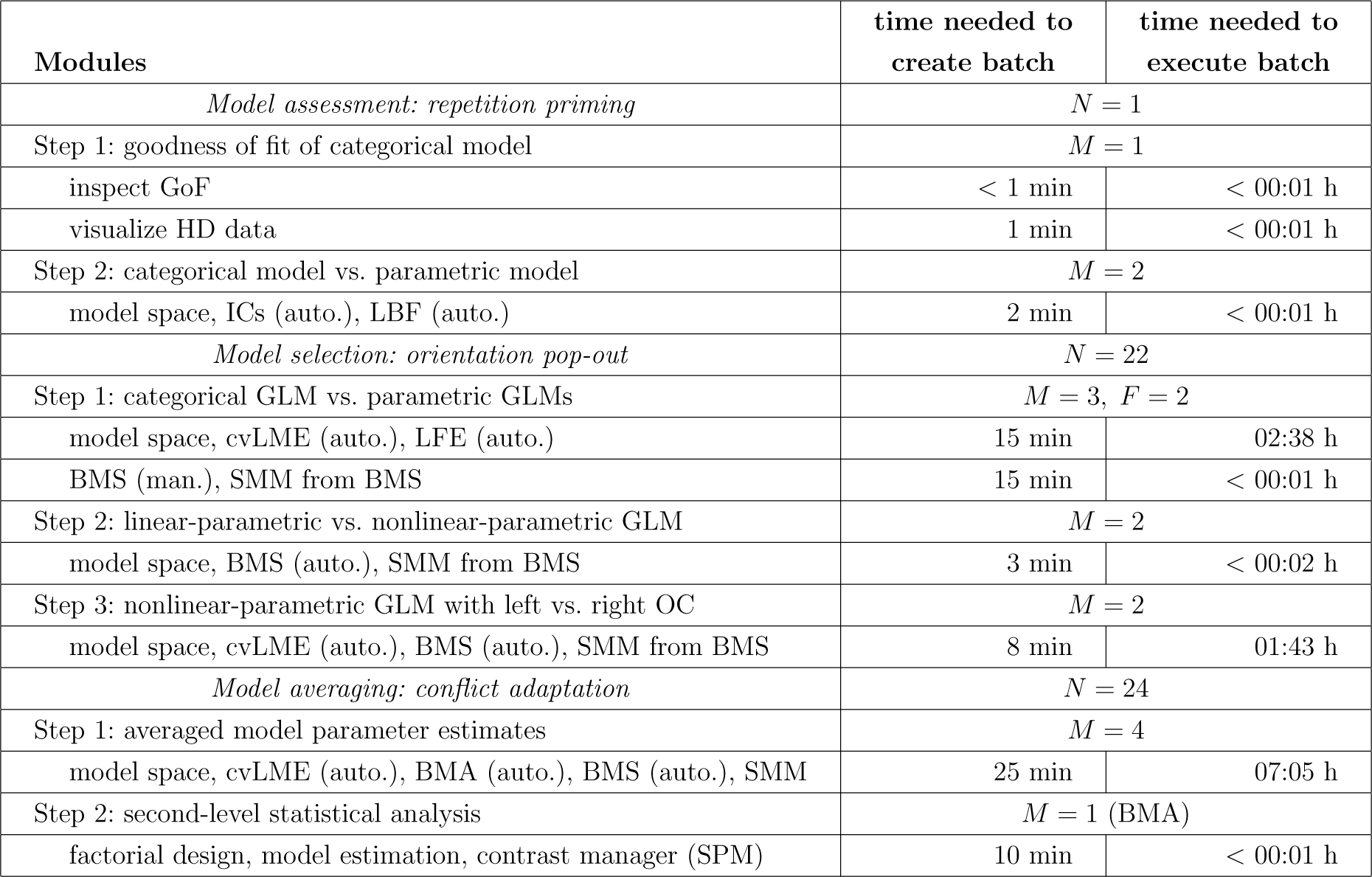
Demonstrations using the MACS toolbox. This table lists processing steps used in first-level model assessment (see Section 4.1) or belonging to cvBMS and cvBMA pipelines (see Sections 4.2 and 4.3) as well as dimensions of the data sets used (*N* = number of subjects, *M* = number of models, *F* = number of families). The second and third column give times required for specifying and running each batch in the SPM batch editor. All computations were performed using SPM12 in MATLAB R2013b running on a 64-bit Windows 7 PC with 16 GB RAM and four hyperthreaded Intel i7 CPU kernels working at 3.40 GHz.

| Modules | time needed to<br>create batch | time needed to<br>execute batch |
| --- | --- | --- |
| *Model assessment: repetition priming* | $N = 1$ | |
| Step 1: goodness of fit of categorical model | $M = 1$ | |
| inspect GoF | < 1 min | < 00:01 h |
| visualize HD data | 1 min | < 00:01 h |
| Step 2: categorical model vs. parametric model | $M = 2$ | |
| model space, ICs (auto.), LBF (auto.) | 2 min | < 00:01 h |
| *Model selection: orientation pop-out* | $N = 22$ | |
| Step 1: categorical GLM vs. parametric GLMs | $M = 3, F = 2$ | |
| model space, cvLME (auto.), LFE (auto.) | 15 min | 02:38 h |
| BMS (man.), SMM from BMS | 15 min | < 00:01 h |
| Step 2: linear-parametric vs. nonlinear-parametric GLM | $M = 2$ | |
| model space, BMS (auto.), SMM from BMS | 3 min | < 00:02 h |
| Step 3: nonlinear-parametric GLM with left vs. right OC | $M = 2$ | |
| model space, cvLME (auto.), BMS (auto.), SMM from BMS | 8 min | 01:43 h |
| *Model averaging: conflict adaptation* | $N = 24$ | |
| Step 1: averaged model parameter estimates | $M = 4$ | |
| model space, cvLME (auto.), BMA (auto.), BMS (auto.), SMM | 25 min | 07:05 h |
| Step 2: second-level statistical analysis | $M = 1$ (BMA) | |
| factorial design, model estimation, contrast manager (SPM) | 10 min | < 00:01 h |

## 5 Discussion

In functional magnetic resonance imaging (fMRI), model quality of general linear models (GLMs) is rarely assessed (Razavi et al., 2003) which may have contributed to the current reproducibility crisis in cognitive neuroscience (Pernet and Poline, 2015). We have introduced an SPM toolbox for model assessment, comparison and selection (MACS) of GLMs for fMRI which implements classical, information-theoretic and Bayesian measures of model quality, most prominently cross-validated Bayesian model selection (cvBMS; Soch et al., 2016) and averaging (cvBMA; Soch et al., 2017).

The key features of our toolbox are (i) *simplicity* – the toolbox is very easy to use and directly operates on SPM.mat files from GLM estimation in SPM; (ii) *generality* – model assessment and selection are not restricted to special model comparisons, but operate on arbitrary design matrices; (iii) *modularity* – toolbox features can be combined into unique processing batches or even appended to existing SPM infrastructure; (iv) *readability* – toolbox code is clearly written for developers and includes extensive comments explaining mathematical background as well as implementational details^7^; and (v) *extendability* – the toolbox can be easily augmented with new modules, e.g. when one wants to employ other variants of first-level model assessment (Penny et al., 2007, eq. 6) or second-level model selection (Rigoux et al., 2014, fig. 8).

The MACS toolbox assists the data analyst in identifying optimal modelling approaches which, in the ideal case, minimizes the distance between the assumed and the actual data generating process, i.e. between the true underlying regularities and the model that is actually employed. In effect, parameter estimates become more precise, e.g. have a lower mean squared error (Soch et al., 2017), and statistical tests become more powerful, i.e. have a higher true positive rate (Soch et al., 2016). On the long run, incorporation of the MACS toolbox into GLM processing pipelines will therefore give rise to fMRI results that are more likely to replicate across studies.

Some of the toolbox features, in particular goodness-of-fit measures and model comparison methods, are more of a diagnostic nature, i.e. to get a qualitative impression of model fit and potentially optimize first-level models. Others, in particular methods of Bayesian model selection and averaging, allow for concrete objective, user-independent modelling decisions. As the diagnostic features are primarily intended to serve exploratory data analysis, we refrain from recommending explicit thresholds for their application which should depend on the specific context.

We want to emphasize that the Bayesian methods in our toolbox such as cvLME calculation do not require a Bayesian estimation of the GLM in SPM. In fact, a model only needs to be specified for model assessment via classical ICs or the Bayesian LME. If a model has been specified but not estimated, the MACS toolbox automatically invokes restricted maximum likelihood (ReML) estimation to obtain the temporal covariance matrix needed for model assessment (Friston et al., 2002). Thanks to a very similar variable structure, the cvLME can also be calculated for GLMs for EEG (Litvak et al., 2011).

Also note that, while MACS is particularly integrated with SPM, it can essentially be used via the MATLAB command line only, with SPM installed by the user, but not being launched at the time of analysis. This is important when users intend to combine our toolbox with other parts of their processing pipelines set up in other analysis packages such as FSL or AFNI – which could e.g. be done using the pipeline manager *Nipype* (Gorgolewski et al., 2011).

Version 1.0 of the MACS toolbox has been uploaded to GitHub in May 2017 (DOI: 10.5281/zenodo.845404). The developers intend to immediately commit bug fixes to this repository (see Section 6). Future developments of the MACS toolbox may include group-level statistical tests for goodness-of-fit measures (Razavi et al., 2003) and information criteria (Vuong, 1989) as well as improved voxel-wise visualization for second-level model selection (cvBMS) results.

## 6 Software Note

The latest version of the MACS toolbox can be downloaded from GitHub (https://github.com/JoramSoch/MACS) and has to be placed as a sub-directory “MACS” into the SPM toolbox folder. Upon starting SPM, batch modules for toolbox features can be assessed by clicking “SPM → Tools → MACS Toolbox” in the SPM batch editor (see Figure 3). The MACS toolbox manual gives detailed instructions how to perform specific analyses.^8^ MACS is optimized for SPM12, but also compatible with SPM8. Consequently, MACS can read all image files (`.nii, .img, .hdr`) that can be read by SPM.

## 7 Acknowledgements

This work was supported by the Bernstein Computational Neuroscience Program of the German Federal Ministry of Education and Research (BMBF grant 01GQ1001C), the Research Training Group “Sensory Computation in Neural Systems” (GRK 1589/1-2), the Collaborative Research Center “Volition and Cognitive Control: Mechanisms, Modulations, Dysfunctions” (SFB 940/1) and the German Research Foundation (DFG grants EXC 257 and KFO 247).

Joram Soch received a Humboldt Research Track Scholarship and an Elsa Neumann Scholarship from the State of Berlin.

The authors would like to thank Carsten Bogler and Achim Meyer for acquiring the fMRI data sets used in this publication.

The authors have no conflict of interest, financial or otherwise, to declare.

## Footnotes

1 Tom Nichols gave a “Requiem for SPMd” on Slide 3 of his talk “Building Confidence in fMRI Results with Model Diagnosis” in the workshop “How Not to Analyze Your Data: A Skeptical Introduction to Modeling Methods” at the 2013 Annual Meeting of the Organization for Human Brain Mapping. A potential reason for the end of support for SPMd could be that it only enables model *diagnosis*, but not actual model *selection* facilitated by the MACS toolbox.

2 The deviance scale, i.e. multiplication by −2, is commonly used in information criteria to remove multiple −1*/*2 factors in the (multivariate normal) log-likelihood function and based on theoretical results obtained with the derivation of the AIC (Burnham and Anderson, 2001).

3 For single-session fMRI data, split-half cross-validation is used and several scans between the two parts are removed to ensure temporal independence (Soch et al., 2016, Fn. 1; Soch et al., 2017, Fn. 1)

4 Note that *m*_*i*_, the optimal model for the *i*-th subject, is a 1 × *M* vector of zeros with a single one at the *j*-th position indicating the true model *j*. Therefore, the model evidence of the *j*-th model is denoted using the *j*-th elementary row vector *e*_*j*_ in the first line of equation (24).

5 An information difference ΔIC < −10 is roughly equivalent to a log Bays factor LBF *>* 5 and thus signals “very strong evidence” according to a widely used scale (Kass and Raftery, 1995).

6 Note that (i) we of course advocate to perform this model comparison using Bayesian model quality measures such as the cvLME rather than information criteria like BIC and (ii) the model comparison performed here is, strictly speaking, not equivalent to the *F* -test in the SPM manual, because the parametric model also excludes regressors that are included in the categorical model, namely experimental condition regressors convolved with the hemodynamic derivatives.

7 Information on any toolbox function (see Figure 1) can be obtained by simply typing `help MACS_fct_name` (e.g. `help MA_cvLME_multi`) into the MATLAB command window after successful MACS toolbox installation (see Section 6). A list of functions with descriptions can be generated using the built-in script `function_index` (resulting in `function_index.xls`).

8 The latest version of the toolbox manual can be found under the following address: https://github.com/JoramSoch/MACS/blob/master/MACSManual/Manual.pdf.

