## Supplementary Materials for "MACS – a new SPM toolbox for model assessment, comparison and selection"

### MACS Manual

Joram Soch, BCCN Berlin

**Toolbox URL:** https://github.com/JoramSoch/MACS

**Support Contact:** Joram Soch ⟨⟩

**Current Version:** MACS V1.1 DOI 10.5281/zenodo.1129353

**Toolbox Paper:** Soch J, Allefeld C (in review). MACS – a new SPM toolbox for model assessment, comparison and selection. *Journal of Neuroscience Methods*, in review, available at *bioRxiv*; DOI: 10.1101/194365.

### Contents


1 How to install the toolbox and read this manual 1

2 How to define a model space of GLMs for fMRI 3

3 How to inspect the goodness of fit for a single model 5

4 How to calculate classical information criteria for a single model 6

5 How to calculate the information difference between two models 7

6 How to calculate the cvLME for a single subject and model 9

7 How to calculate the cvLME for multiple subjects or models 10

8 How to calculate posterior probabilities from cvLMEs 11

9 How to calculate log family evidences from cvLMEs 12

10 How to calculate log Bayes factors from cvLMEs 13

11 How to perform cross-validated Bayesian model selection 14

12 How to perform cross-validated Bayesian model averaging 16

13 How to visualize first-level model averaging results 18

14 How to visualize second-level model selection results 19

15 How to extend the toolbox with your own modules 21

### 1 How to install the toolbox and read this manual

**Purpose:** This is a manual for **MACS** (pronounced as "Max"), an SPM toolbox for model assessment, comparison and selection (MACS) of general linear models (GLMs) applied to functional magnetic resonance imaging (fMRI) data using Statistical Parametric Mapping (SPM). MACS includes classical, information-theoretic and Bayesian methods of model assessment previously applied to GLMs for fMRI as well as recent methodological developments of model selection (Soch et al., 2016) and model averaging (Soch et al., 2017) in fMRI data analysis (Soch & Allefeld, in review).

**Requirements:** To use the MACS toolbox, you need SPM8 or SPM12 running in MATLAB R2008a or later running on any operating system.

**Procedure:** To install the MACS toolbox, proceed as follows:

- Go to the GitHub repository of the MACS toolbox (see URL given on title page).
- Download the toolbox by clicking "Clone or download → Download ZIP".
- Unpack the ZIP file and rename the folder from "MACS-master" to "MACS".
- Place the folder into SPM's toolbox directory. This directory can be retrieved by typing into the MATLAB command window: `strcat(spm('dir'),'/toolbox/')`.
- Launch SPM by typing `spm fmri` into the MATLAB command window.
- Open the SPM batch editor by clicking "Batch" in the SPM menu window.

**Consequences:** Upon successful installation of the toolbox, its modules can be accessed by clicking "SPM → Tools → MACS Toolbox" (see Figure 1).

**Remark:** When reading this manual, you are not meant to go from first to last page, but instead choose a specific analysis to be performed and search for the corresponding section in the manual. Each section is subdivided into subsesctions (*Purpose – Requirements – Procedure – Consequences*) and is written in the form of a cooking recipe, so that you can follow step-by-step instructions to perform a specific analysis. If you are unsure which analysis you want to perform or about where to start, we recommend to read the manual in the following order: Sections 3 – 6 – 2 – 7 – 11 – 14 – 12 – 13.

**Documentation:** Further information on the toolbox can be found in two theoretical papers (Soch et al., 2016; Soch et al., 2017) as well as a toolbox paper (Soch & Allefeld, in review) and a technical report (Soch, in prep.) which you are also welcome to cite when using the toolbox:

- Soch J, Haynes JD, Allefeld C (2016). How to avoid mismodelling in GLM-based fMRI data analysis: cross-validated Bayesian model selection. *NeuroImage*, vol. 141, pp. 469-489; DOI: 10.1016/j.neuroimage.2016.07.047.
- Soch J, Meyer AP, Haynes JD, Allefeld C (2017). How to improve parameter estimates in GLM-based fMRI data analysis: cross-validated Bayesian model averaging. *NeuroImage*, vol. 158, pp. 186-195; DOI: 10.1016/j.neuroimage.2017.06.056.
- Soch J, Allefeld C (in review). MACS – a new SPM toolbox for model assessment, comparison and selection. *Journal of Neuroscience Methods*, in review, available at *bioRxiv*; DOI: 10.1101/194365.
- Soch J (in prep.). cvBMS and cvBMA: filling in the gaps. Technical Report, in preparation, will be available at *arXiv*.

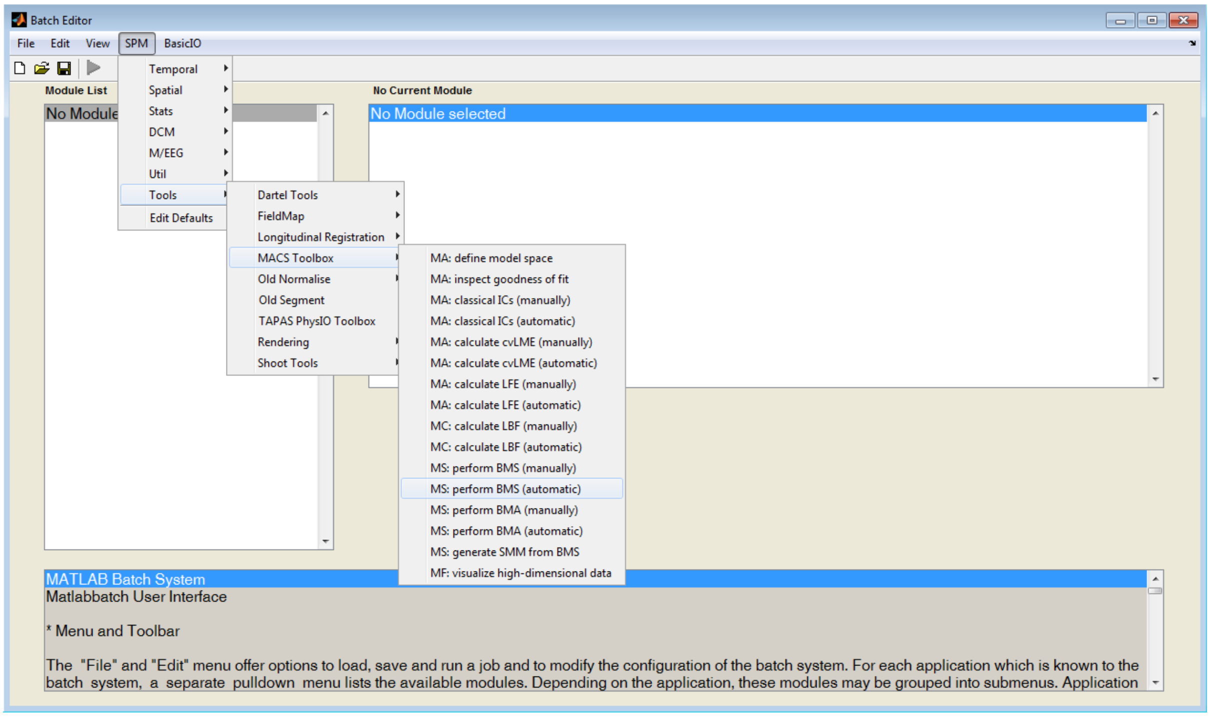


**Figure 1.** MACS toolbox modules in the SPM batch editor.

### 2 How to define a model space of GLMs for fMRI

**Purpose:** Defining a model space enables you to use the full power of the automatic processing mode provided by the MACS toolbox. In short, whereas manual processing requires you to enter filepaths of the models in each individual processing step, they only need to be specified once in automatic processing. In this manual, Sections 3–6 will use the manual processing mode while Sections 7–12 will use the automatic processing mode and thus rest on defining a model space.

**Requirements:** This module requires that several models ($M$ = number of models) have been specified for several subjects ($N$ = number of subjects) and the `SPM.mat` files for all these models are available.

**Procedure:** Defining a model space proceeds as follows:

- Open the SPM batch editor by clicking "Batch" in the SPM menu window.
- Select the MACS toolbox module for model space definition by clicking "SPM → Tools → MACS Toolbox → MA: define model space".
- Choose a model space directory by clicking "Directory → Specify". This will be the directory into which the model space file `MS.mat` will be saved and where second-level analysis results will be written, e.g. when performing group-level Bayesian model selection (see e.g. Section 11).
- Create a new subject by clicking "Select SPM.mats → New: Subject".
- Create a new model by clicking "Subject → New: Model".
- Select the `SPM.mat` belonging to this model by clicking "Model → Specify". This model has to be specified in SPM, but does not necessarily need to be estimated.
- Repeat the last three steps for all models applied to all subjects. The number of models and their order of specification has to be the same for all subjects.
- Create a new model label by clicking "Enter GLM names → New: Name".
- Enter the name of this model by clicking "Name → Specify".
- Repeat the last two steps for all models in the model space. The order of specification has to match the order of the `SPM.mat` files within each subject.

- Save the batch under the filename `batch.mat` into the model space directory. This is recommended for easier setup of later analyses, e.g. when initializing the model space with a different model space directory.
- Run the batch by clicking the green triangle.

**Consequences:** This module results in an `MS.mat` file saved into the model space directory specified above. This file contains a single struct variable `MS` with the fields `swd`, a string specifying the model space directory; `SPMs`, an $N \times M$ cell array of `SPM.mat` filepaths; and `GLMs`, an $1 \times M$ cell array of GLM names (see Figure 2). Such an `MS.mat` file can be used as an input parameter by any MACS toolbox module belonging to the automatic processing stream (see e.g. Section 7).

**Remark:** When the number of subjects or the number of models is very large, even the automatic processing mode relying on model space definition can be tedious. At least for programmers, a simpflification is available as a template script in the MACS toolbox. Simply open the file `MS_mat_pipeline.m` from the toolbox sub-directory `MACS_Pipelines` and edit subject IDs, model labels as well as output directory. This script works, if your data are organized in a subject-model hierarchy, and writes the `MS.mat` file as well as an editable `batch.mat` file into the specified directory.

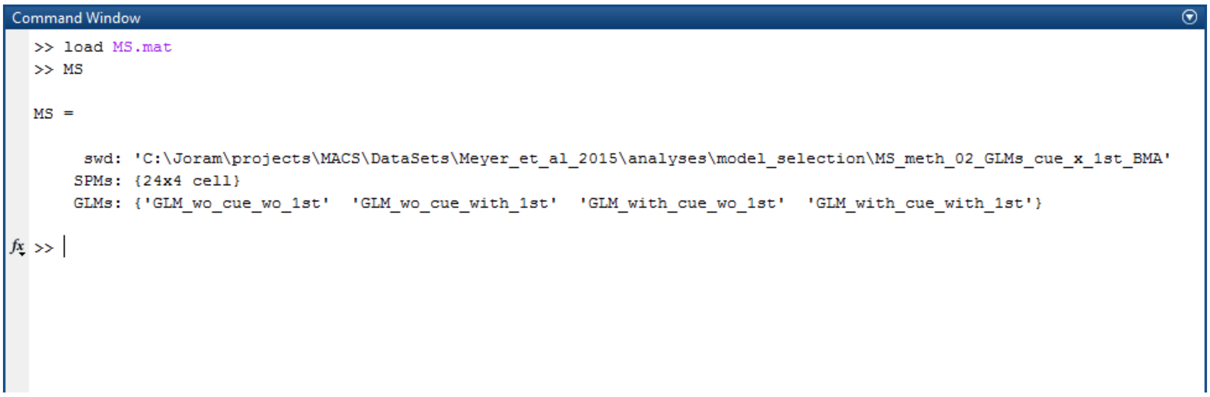


**Figure 2.** Structure of a model space variable in a model space file.

### 3 How to inspect the goodness of fit for a single model

**Purpose:** Assessing the goodness of fit for a single GLM allows a visual inspection of how well a particular model fits the measured data. Additionally, it will give rise to metrics that allow to quantify and compare the goodness of fit.

**Requirements:** This module requires that one model has been estimated in SPM, with its `SPM.mat` file being available on disk.

**Procedure:** Inspecting the goodness of fit proceeds as follows:

- Open the SPM batch editor by clicking "Batch" in the SPM menu window.
- Select the MACS toolbox module for goodness-of-fit inspection by clicking "SPM → Tools → MACS Toolbox → MA: inspect goodness of fit".
- Select the model by clicking "Select SPM.mat → Specify". This model needs to be specified and estimated in SPM.
- Select the plotting type by highlighting "Type of plot" and clicking "line graph" or "scatter plot". The suggested and default option is "line graph".
- Save the batch under the filename `MA_inspect_GoF.mat` into the GLM directory.
- Run the batch by clicking the green triangle.

**Consequences:** This module results in an interactive plot in the SPM graphics window. The lower panel will show an anatomical image highlighting the top 5% (good fit, in yellow color) and the bottom 5% (bad fit, in red color) of the voxels in terms of "R-squared", a measure for model fit. Upon browsing through through the anatomical image, the upper panel will adapt instantly and show the measured signal against the predicted signal as well as goodness-of-fit measures for the current voxel.

Furthermore, several images will be written into the GLM directory: `MA_GoF_R2.nii`, the coefficient of determination $R^2$; `MA_GoF_R2_adj.nii`, the adjusted coefficient of determination $R^2_{\mathrm{adj}}$; `MA_GoF_SNR_mf.nii`, the model-free signal-to-noise ratio $\mathrm{SNR}_{\mathrm{mf}}$; and `MA_GoF_SNR_mb.nii`, the model-based signal-to-noise ratio $\mathrm{SNR}_{\mathrm{mb}}$.

### 4 How to calculate classical information criteria for a single model

**Purpose:** Classical information criteria (ICs) are model selection criteria that are based on the maximum log-likelihood (MLL), as a measure of model accuracy, and some function of the number of data points $n$ and the number of model parameters $p$, as a measure of model complexity. Classical ICs allow to quantify the quality of a model and can be directly compared between models.

**Requirements:** This module requires that one model has been estimated in SPM, with its `SPM.mat` file being available on disk.

**Procedure:** Computing classical ICs proceeds as follows:

- Open the SPM batch editor by clicking "Batch" in the SPM menu window.
- Select the MACS toolbox module for classical information criteria by clicking "SPM → Tools → MACS Toolbox → MA: classical ICs (manually)".
- Select the model by clicking "Select SPM.mat → Specify". This model needs to be specified and estimated in SPM. Note that the classical IC module can also be directly appended to existing SPM batch jobs. If this is the case, select the model by clicking "Select SPM.mat → Dependency" and then e.g. "fMRI model specification: SPM.mat file → OK" or "Model estimation: SPM.mat file → OK".
- Select the model quality criterion by highlighting "Information criterion" and clicking one of the criteria. The two most common ones are "AIC" and "BIC".
- Save the batch under the filename `MA_ICs_XIC.mat` into the GLM directory where `XIC` is to be replaced by the selected information criterion.
- Run the batch by clicking the green triangle.

**Consequences:** This module results in an image called `MA_ICs_XIC.nii` in the GLM directory where `XIC` will be the selected information criterion. When this has been done for two models, an information difference can be calculated (see Section 5).

### 5 How to calculate the information difference between two models

**Purpose:** The information difference ($\Delta$IC) is simply the difference between the information criteria for two models (see Section 4). It can be used (i) to find out which model better explains the data and (ii) to assess the amount of evidence that there is for either of the models, e.g. via posterior probabilities.

**Requirements:** These operations require that two models for the same subject have been estimated in SPM, with their `SPM.mat` files being available on disk.

**Procedure:** Computing an information difference proceeds as follows:

- Set up a batch that calculates the desired information criterion for the first model (see Section 4), but don't save and run it.
- Create a new module by right-clicking "MA: classical ICs (manually)" and then left-clicking "Replicate Module".
- In this module, select the second model by clicking "Select SPM.mat → Specify".
- Select the MACS toolbox module for log Bayes factor by clicking "SPM → Tools → MACS Toolbox → MC: calculate LBF (manually)". Although this module is primarily designed for calculating log Bayes factors (LBFs), i.e. differences between log model evidences (LMEs), it can in principle be used to calculate the difference between any two model quantities, e.g. also information criteria.
- Choose an output directory by clicking "Directory → Specify". This will be the directory where the information difference images will be saved.
- Create a new subject by clicking "Select LME maps → New: Subject".
- Create a new model by clicking "Subject → New: Model".
- Select the first model by clicking "Model → Dependency" and then selecting the output from the first module. This creates a reference to the information criterion image file that will be calculated when executing the first module belonging to the first model.
- Repeat the last two steps in order to select the second model.
- Create a new model label by clicking "Enter GLM names → New: Name".
- Enter the name of the first model by clicking "Name → Specify".
- Repeat the last two steps in order to name the second model.

- Save the batch under the filename `design.mat` into the output directory.
- Run the batch by clicking the green triangle.

**Consequences:** These operations result in several images written into the output directory (see Figure 3), among them `LBF_GLM_1_vs_GLM_2_sub1.nii`, the information difference in favor of the first model, i.e. $\Delta\mathrm{IC}_{12} = \mathrm{IC}(m_1) - \mathrm{IC}(m_2)$; and `LBF_GLM_2_vs_GLM_1_sub1.nii`, the information difference in favor of the second model, i.e. $\Delta\mathrm{IC}_{21} = \mathrm{IC}(m_2) - \mathrm{IC}(m_1)$; the other images can be ignored.

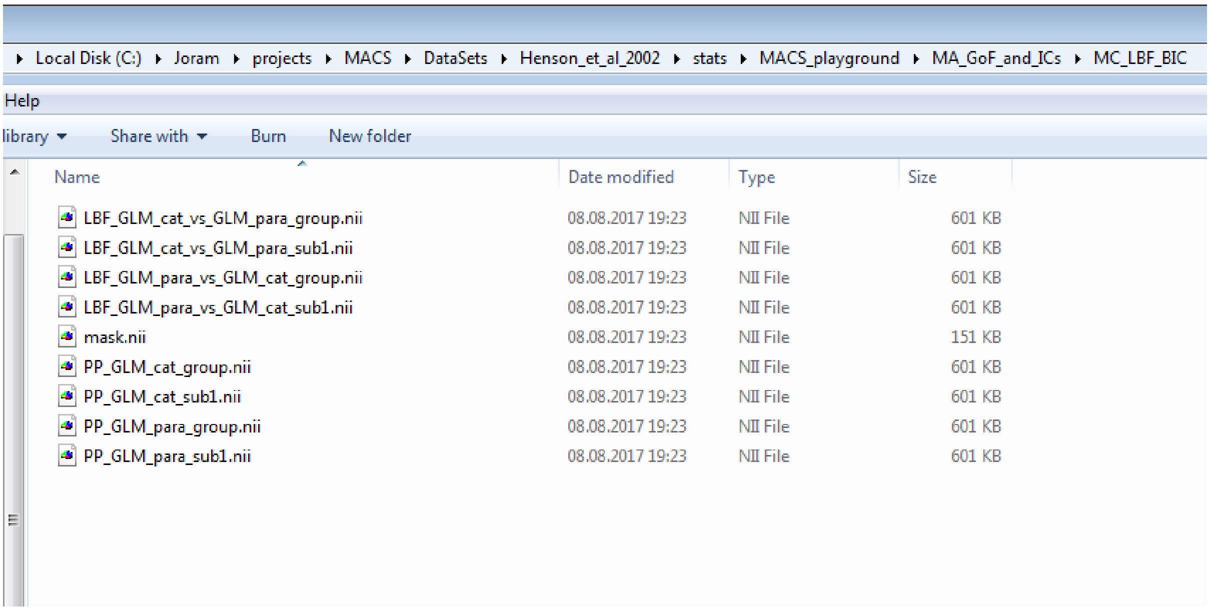


**Figure 3.** File output from application of the log Bayes factor module.

### 6 How to calculate the cvLME for a single subject and model

**Purpose:** The cross-validated log model evidence (cvLME) is a model selection criterion that combines cross-validation with the Bayesian marginal likelihood and underlies cross-validated Bayesian model selection (cvBMS) and averaging (cvBMA).

**Requirements:** This module requires that one model has been specified in SPM, with its `SPM.mat` file being available on disk.

**Procedure:** Computing the cvLME proceeds as follows:

- Open the SPM batch editor by clicking "Batch" in the SPM menu window.
- Select the MACS toolbox module for cross-validated log model evidences by clicking "SPM → Tools → MACS Toolbox → MA: calculate cvLME (manually)".
- Select the model by clicking "Select SPM.mat → Specify". This model needs to be specified and/or estimated in SPM. Note that the cvLME module can also be directly appended to existing SPM batch jobs. If this is the case, select the model by clicking "Select SPM.mat → Dependency" and then e.g. "fMRI model specification: SPM.mat file → OK" or "Model estimation: SPM.mat file → OK".
- Choose whether model accuracy and model complexity are calculated by clicking "Accuracy & Complexity → Yes/No". Generally, if you don't intend to look at them in detail, it is recommended that this option is left at the default "No" in order save processing time and disk space.
- Save the batch under the filename `MA_cvLME.mat` into the GLM directory.
- Run the batch by clicking the green triangle.

**Consequences:** This module results in an image called `MA_cvLME.nii` in the GLM directory which contains the voxel-wise cross-validated log model evidence for this model. Furthermore, a number of images `MA_cvLME_SX.nii` where `X` ranges from 1 to the number of fMRI recording sessions $S$ will be saved that contain voxel-wise out-of-sample log model evidences for each session. If the selected first-level model is a single-session design, such that split-half cross-validation is used, these images will be called `MA_cvLME_P1.nii` and `MA_cvLME_P2.nii`. Usually, only the cross-validated and not the out-of-sample evidences will be needed and can facilitate a lot of further operations (see Section 7).

### 7 How to calculate the cvLME for multiple subjects or models

**Purpose:** Computing the cross-validated log model evidence (cvLME) for multiple subjects or models automatically is advantageous when the number of subjects (studied cohort) or the number of models (model space) is large.

**Requirements:** This module requires a model space file called `MS.mat` (see Section 2) listing a number of models for a number of subjects.

**Procedure:** Computing the cvLME proceeds as follows:

- Open the SPM batch editor by clicking "Batch" in the SPM menu window.
- Select the MACS toolbox module for cross-validated log model evidences by clicking "SPM → Tools → MACS Toolbox → MA: calculate cvLME (automatic)".
- Select the model space by clicking "Select MS.mat → Specify".
- Run the batch by clicking the green triangle.
- Really, this is all? Yes, this is it!

**Consequences:** This module results in an image called `MA_cvLME.nii` for the cross-validated and several images `MA_cvLME_SX.nii` for the out-of-sample log model evidences (see Section 6) in the directory of each GLM that is in the model space specified by the `MS.mat` file. These cvLME maps can then be further processed into posterior probabilities (see Section 8), log family evidences (see Section 9), log Bayes factors (see Section 10) or used for cross-validated Bayesian model selection (see Section 11) and cross-validated Bayesian model averaging (see Section 12).

### 8 How to calculate posterior probabilities from cvLMEs

**Purpose:** Because the cvLME is a relative measure of model quality, its absolute value has no direct interpretation. However, posterior probabilities (PP) can be used to quantify the evidence in favor of each model. This section demonstrates how to calculate posterior model probabilities for a single subject.

**Requirements:** This module requires that several models for the same subject have been assessed using the cvLME (see Section 6 or 7), such that a cvLME map `MA_cvLME.nii` exists in the directory of each.

**Procedure:** Computing posterior probabilities proceeds as follows:

- Go to the folder belonging to this subject, i.e. where the model sub-directories are located, and create a new sub-directory, e.g. `MS_BMS_subject`. This is the output directory into which analysis results will be saved.
- Set up a batch that defines a model space (see Section 2) which only includes the models from this particular subject and uses the output folder as the model space directory. Don't save and run this batch!
- Select the MACS toolbox module for group-level Bayesian model selection by clicking "SPM → Tools → MACS Toolbox → MS: perform BMS (automatic)".
- Select the model space by clicking "Select MS.mat → Dependency" and then "MA: define model space: model space (MS.mat file) → OK".
- Switch the estimation algorithm by clicking "Inference method → Fixed Effects (FFX)".
- Save the batch under the filename `design.mat` into the output directory.
- Run the batch by clicking the green triangle.

**Consequences:** This module results in an image called `GLM_XYZ_model_PPM.nii` for each model defined in the model space where `GLM_XYZ` stands for the name given to this model. This posterior probability map (PPM) contains the voxel-wise posterior probabilities of this model relative to the cvLMEs from the whole model space.

### 9 How to calculate log family evidences from cvLMEs

**Purpose:** When some models in a model space are more similar to each other than they are to another model, model families need to be formed for proper model selection and to avoid an unfair comparison. This can be achieved by calculating log family evidences (LFEs) from log model evidences (LMEs).

**Requirements:** This module requires that several models for one or more subjects have been assessed using the cvLME (see Section 6 or 7), such that a cvLME map `MA_cvLME.nii` exists in the directory of each.

**Procedure:** Computing log family evidences proceeds as follows:

- Set up a batch that defines a model space (see Section 2) including all models from all subjects, but don't save and run this batch!
- Select the MACS toolbox module for log family evidence calculation by clicking "SPM → Tools → MACS Toolbox → MA: calculate LFE (automatic)".
- Select the model space by clicking "Select MS.mat → Dependency" and then "MA: define model space: model space (MS.mat file) → OK".
- Specify the family affiliation by clicking "Family vector → Specify". This has to be a $1 \times M$ vector ($M$ = number of models) where the number $j$ in the $i$-th position indicates that the $i$-th model belongs to family $j$. For example, if you have 10 models belonging to 4 families, the family vector could be `[1 1 1 2 2 2 3 3 4 4]`.
- Create a new family label by clicking "Family names → New: Name".
- Enter the name of this model family by clicking "Name → Specify".
- Repeat the last two steps for all model families. The order of specification has to match the numbers in the family vector and it is required that the numer of family names is equal to the highest number in the family vector.
- Save the batch under the filename `design.mat` into the model space directory.
- Run the batch by clicking the green triangle.

**Consequences:** For each subject in the model space, this module will create an output directory called `MA_LFE_uniform` in the folder where the first model's sub-directory is located. Within this folder, there will be an image called `GLMs_XYZ_LFE.nii` for each model family where `GLMs_XYZ` stands for the name given to this family. These log family evidence (LFE) maps can then be used for further statistics. Generally speaking, all analyses that operate on cvLME maps, can also operate on LFE maps.

### 10 How to calculate log Bayes factors from cvLMEs

**Purpose:** When only comparing two models, one can compute log Bayes factors (LBFs) from log model evidences (LMEs). An LBF is nothing else than a difference between two LMEs, assumes a uniform prior over models and can be subjected to certain scales of interpretation (see below).

**Requirements:** This module requires that two models for one or more subjects have been assessed using the cvLME (see Section 6 or 7), such that a cvLME map `MA_cvLME.nii` exists in the directory of each.

**Procedure:** Computing log Bayes factors proceeds as follows:

- Create a new directory, e.g. `MS_LBF_group`, at an arbitrary location, preferably somewhere in your second-level results folder. This is the output directory into which analysis results will be saved.
- Set up a batch that defines a model space (see Section 2) including both models from all subjects and using the output folder as the model space directory. Don't save and run this batch!
- Select the MACS toolbox module for log Bayes factor calcualtion by clicking "SPM → Tools → MACS Toolbox → MC: calculate LBF (automatic)".
- Select the model space by clicking "Select MS.mat → Dependency" and then "MA: define model space: model space (MS.mat file) → OK".
- Save the batch under the filename `design.mat` into the output directory.
- Run the batch by clicking the green triangle.

**Consequences:** For each subject and model in the model space, this module will write an image called `LBF_GLM_1_vs_GLM_2_subX.nii` giving the voxel-wise log Bayes factor (LBF) and `PP_GLM_1_subX.nii` giving the voxel-wise posterior probability (PP) into the output directory where `X` ranges from 1 to the number subjects $N$ and `GLM_1/2` are the names of the two models. Additionally, such LBF and PP images with filenames ending on `_group.nii` instead of `_subX.nii` will be saved for the entire group of subjects which is based on the log group Bayes factor (LGBF) method.

### 11 How to perform cross-validated Bayesian model selection

**Purpose:** Cross-validated Bayesian model selection (cvBMS; Soch et al., 2016) is a principled approach to avoid mismodelling in GLM-based fMRI data analysis, a method that allows to choose the best GLM for a given group-level fMRI data set. It consists in (i) first-level model assessment using the cross-validated log model evidence (cvLME), (ii) second-level model inference using random-effects Bayesian model selection (RFX BMS) and (iii) voxel-wise model selection by best-performing model identification.

**Requirements:** These operations require that several models for several subjects have been specified in SPM, with their `SPM.mat` files available on disk.

**Procedure:** To perform cvBMS, proceed as follows:

- Create a new directory, e.g. `MS_cvBMS`, at an arbitrary location, preferably somewhere in your second-level results folder. This is the output directory into which analysis results will be saved.
- Set up a batch that defines a model space (see Section 2) which includes all models from all subjects and uses the output folder as the model space directory. Don't save and run this batch!
- Select the MACS toolbox module for cross-validated log model evidences by clicking "SPM → Tools → MACS Toolbox → MA: calculate cvLME (automatic)".
- Select the model space by clicking "Select MS.mat → Dependency" and then "MA: define model space: model space (MS.mat file) → OK".
- Select the MACS toolbox module for group-level Bayesian model selection by clicking "SPM → Tools → MACS Toolbox → MS: perform BMS (automatic)".
- Select the model space by clicking "Select MS.mat → Dependency" and then "MA: define model space: model space (MS.mat file) → OK".
- Select the MACS toolbox module for selected-model maps after BMS by clicking "SPM → Tools → MACS Toolbox → MS: generate SMM from BMS".
- Select model selection results by clicking "Select BMS.mat → Dependency" and then "MS: perform BMS (automatic): BMS results (BMS.mat file) → OK".

- Save the batch under the filename `design.mat` into the output directory.
- Run the batch by clicking the green triangle.

**Consequences:** For each model in the model space, these operations will write the following images into the output directory: `GLM_XYZ_model_alpha.nii`, voxel-wise concentration parameters of the posterior Dirichlet distribution that comes out of the RFX BMS; `GLM_XYZ_model_EF.nii`, the expected frequency (EF) of each model, i.e. the posterior mean of model frequencies; `GLM_XYZ_model_LF.nii`, the likeliest frequency (EF) of each model, i.e. the posterior mdoe of model frequencies; and, if the attribute "Exceedance probabilities" in the module "MS: perform BMS (automatic)" was set to "Yes", `GLM_XYZ_model_EP.nii`, the exceedance probability of each model, i.e. the posterior probability that this model is more frequent (or, "more often used") than any other model in the model space. Generally, the most interesting thing to look at are likeliest frequencies which can be interpreted as the proportion of subjects in which the corresponding model is the optimal model or best explanation of the measured data.

Additionally, there will be sub-directory called `MS_SMM_BMS_10` which contains the following images: `MS_SMM_map_pos_X_mod_Y_GLM_XYZ.nii`, where `GLM_XYZ` is the name of the model placed at the `Y`-th position when defining the model space and ranking on `X`-th place when ordering by the number of voxels in which models are selected according to estimated model frequency. For example, `MS_SMM_map_pos_2_mod_8_GLM_B4.nii` would be the selected-model map (SMM) of `GLM_A4`, the second model in the model space (`mod_8`) and the winning model in second-most of the voxels (pos_2). SMMs are likeliest frequency maps, but only show those voxels where the corresponding model is the optimal model. For further information, type `help MS_SMM_BMS` into the command window.

**Remark:** The pipeline described here is also available as a template batch in the MACS toolbox. Simply start the SPM batch editor and open the file `cvBMS_template.mat` from the toolbox sub-directory `MACS_Pipelines`.

### 12 How to perform cross-validated Bayesian model averaging

**Purpose:** Cross-validated Bayesian model averaging (cvBMA; Soch et al., 2017) is a principled approach to improve parameter estimates in GLM-based fMRI data analysis, a technique that allows to improve parameter estimates by combining several GLMs for a given subject-level fMRI data set. It consists in (i) first-level model assessment using the cross-validated log model evidence (cvLME) and (ii) first-level model combination using standard Bayesian model averaging (BMA).

**Requirements:** These operations require that several models for several subjects have been specified in SPM, with their `SPM.mat` files available on disk.

**Procedure:** To perform cvBMA, proceed as follows:

- Set up a batch that defines a model space (see Section 2) including all models from all subjects, but don't save and run this batch!
- Select the MACS toolbox module for cross-validated log model evidences by clicking "SPM → Tools → MACS Toolbox → MA: calculate cvLME (automatic)".
- Select the model space by clicking "Select MS.mat → Dependency" and then "MA: define model space: model space (MS.mat file) → OK".
- Select the MACS toolbox module for group-level Bayesian model selection by clicking "SPM → Tools → MACS Toolbox → MS: perform BMA (automatic)".
- Select the model space by clicking "Select MS.mat → Dependency" and then "MA: define model space: model space (MS.mat file) → OK".
- Specify the parameter assignment by clicking "Parameter matrix → Specify". This has to be an $M \times P$ matrix ($M$ = number of models, $P$ = number of parameters) which indicates which regressor (column) in which GLM (row) belongs to which common model parameter. For example, if you have three models and four parameters to be averaged, the parameter matrix could be `[1 2 3 4; 1 3 5 7; 1 4 7 10]`. If a model does not include certain model parameters, this would be indicated by entering zeros, e.g. `[1 2 3 4; 1 2 0 0]`. If the parameters have the same indices in all models, it can also be a $1 \times P$ vector, e.g. `[1 2 3 4]`. For more information, read the help text that appears in the batch editor when clicking on "Parameter matrix".

- Choose an analysis title by clicking "Analysis name → Specify". This will allow to distinguish your analysis from others in the same model space, e.g. when only using a subset of the models for model averaging. If you only intend to do one BMA analysis, just leave this parameter at its default value.
- Save the batch under the filename `design.mat` into the model space directory.
- Run the batch by clicking the green triangle.

**Consequences:** For each subject in the model space, this module will create an output directory called `MS_BMA_subject_ba_BMA1` in the folder with the first model's sub-directory where `BMA1` is the name given to the analysis. Within this folder, there will be an image for each parameter called `beta_NNNN_BMA.nii` with the averaged model parameter estimates where `NNNN` is a four-digit index, e.g. `0005`, that is based on the indices in the first row of the parameter matrix, i.e. equivalent to the regressor order from the first model. These averaged parameter (BMA) maps can then be used for further statistics, e.g. second-level tests across subjects. Generally speaking, all analyses that operate on GLM beta maps, can also operate on BMA beta maps.

**Remark:** The pipeline described here is also available as a template batch in the MACS toolbox. Simply start the SPM batch editor and open the file `cvBMA_template.mat` from the toolbox sub-directory `MACS_Pipelines`.

### 13 How to visualize first-level model averaging results

**Purpose:** This section introduces you how to get an impression how an averaged model parameter estimate obtained from cross-validated Bayesian model averaging differs from the individual model's parameter estimates.

**Requirements:** This module requires that cross-validated Bayesian model averaging (cvBMA; see Section 12) has been performed.

**Procedure:** Visualizing cvBMA results proceeds as follows:

- Open the SPM batch editor by clicking "Batch" in the SPM menu window.
- Select the MACS toolbox module for high-dimensional data visualization by clicking "SPM → Tools → MACS Toolbox → MF: visualize high-dimensional data".
- Choose a particular subject from your model space and change MATLAB's current directory to this subject's directory. Then go back to the batch editor.
- Specify the input images by clicking "Input images → Specify". For a particular regressor whose coefficients you have averaged, choose this regressor's beta map from each model's sub-directory as well as this regressor's BMA map `beta_NNNN_BMA.nii` from the sub-directory `MS_BMA_subject_ba_BMA1`.
- Click "Overlay image → Specify" and choose the same BMA map.
- Click "Overlay threshold → Specify" and enter "$>0$" (without quotes).
- Click "Title → Specify" and enter a figure name, e.g. "cvBMA visualization".
- Save the batch under the filename `beta_NNNN_BMA.mat` into the BMA directory.
- Run the batch by clicking the green triangle.

**Consequences:** This module results in an interactive plot in the SPM graphics window. The lower panel will show an anatomical image highlighting the voxels in which the averaged estimate is positive (or negative, if you enter "$< 0$" as the overlay threshold). Upon browsing through through the anatomical image, the upper panel will adapt instantly and show a bar plot of model-wise as well as averaged parameters for the regressor you have chosen. The right-most bar representing the BMA estimate should always be between the minimum and maximum of the other bars.

**Remark:** In the future, this feature will be automatized and have its own module.

### 14 How to visualize second-level model selection results

**Purpose:** This section introduces you how to visualize model frequencies obtained from cross-validated Bayesian model selection.

**Requirements:** This module requires that cross-validated Bayesian model selection (cvBMS; see Section 11) has been performed.

**Procedure:** Visualizing cvBMS results proceeds as follows:

- Open the SPM batch editor by clicking "Batch" in the SPM menu window.
- Select the MACS toolbox module for high-dimensional data visualization by clicking "SPM → Tools → MACS Toolbox → MF: visualize high-dimensional data".
- Search where the analysis results have been saved and change MATLAB's current directory to this cvBMS results directory. Then go back to the batch editor.
- Specify the input images by clicking "Input images → Specify". Choose likeliest frequency maps ending on `_LFM.nii` for all models that were in your analysis.
- Specify the overlay image by clicking "Overlay image → Specify". Choose the selected model map (SMM), located in the sub-directory `MS_SMM_BMS_10` of the cvBMS results folder, from one of the models that you are particularly interested in, e.g. representing the hypothesis that you want to support using your experiment.
- Click "Overlay threshold → Specify" and enter "$>0$" (without quotes).
- Click "X-Axis Ticks → Specify" and enter "`cellstr(num2str([1:M]'))'`". Here, `M` has to be replaced by the number of models in your model space.
- Click "Y-Axis Limits → Specify" and enter "$[0,1]$".
- Click "Title → Specify" and enter a figure name, e.g. "cvBMS visualization".
- Save the batch under the filename `BMS_LFM.mat` into the cvBMS results directory.
- Run the batch by clicking the green triangle.

**Consequences:** This module results in an interactive plot in the SPM graphics window (see Figure 4). The lower panel will show an anatomical image highlighting the voxels in which the model that you are interested in is selected as the optimal model. Upon browsing through through the anatomical image, the upper panel will adapt instantly and show a bar plot of estimated model frequencies.

**Remark:** In the future, this feature will be automatized and have its own module.

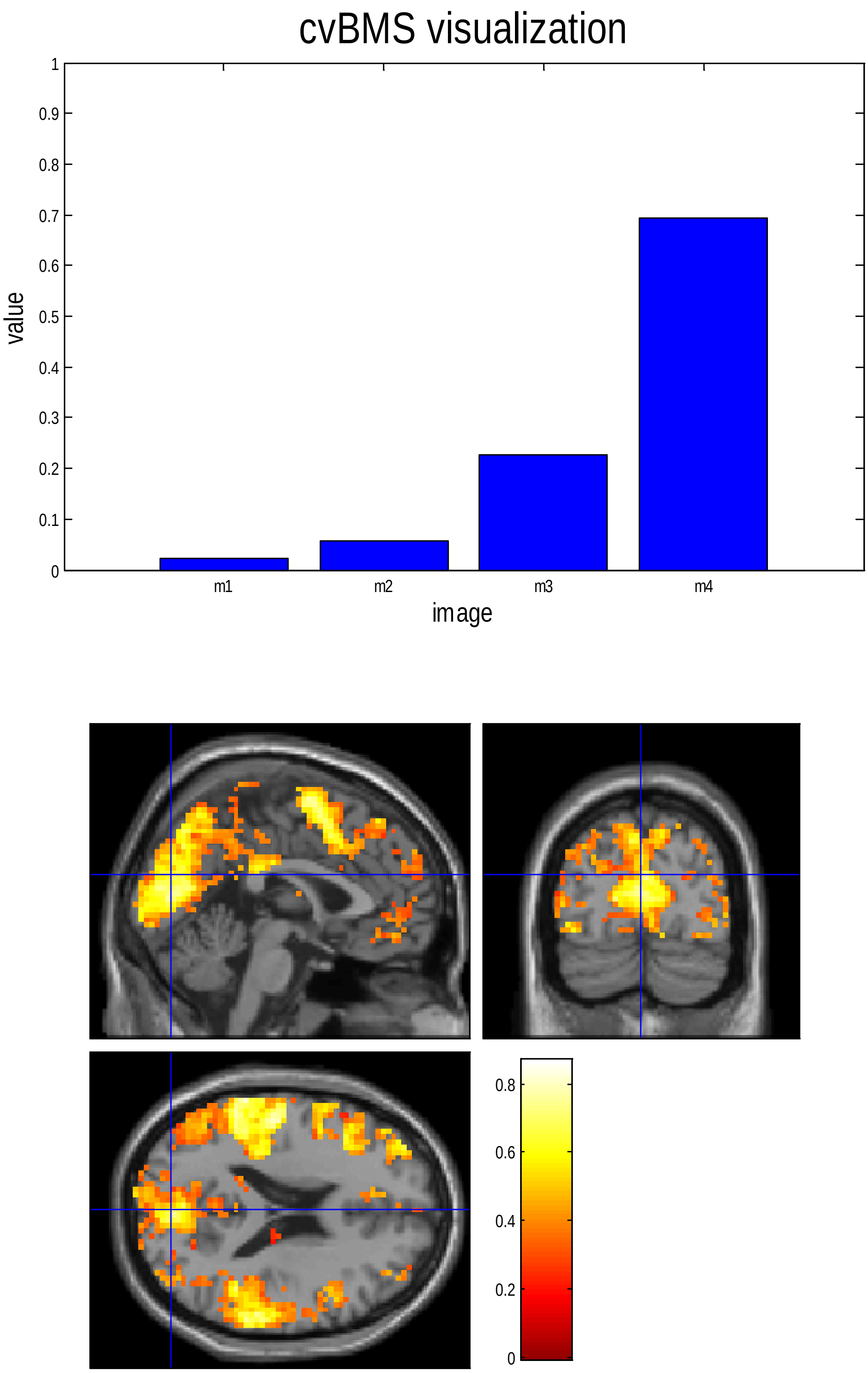


**Figure 4.** Graphical output from second-level model selection results visualization.

### 15 How to extend the toolbox with your own modules

The MACS toolbox can be easily augmented with your own modules, e.g. when you want to employ other variants of first-level model assessment or second-level model selection. Generally, such an extension has to consist of three components:

- a *mathematical function* that implements the desired method;
- an *interface function* that applies the method to GLMs in SPM;
- a *batch function*, if you want this method to be available in the batch editor.

In MACS, references from second-level analyses to first-level images are managed by simply saving the image headers into the `SPM.mat` file as fields of the struct variable `SPM.MACS`. Consequently, if you want to implement a new first-level criterion called ABC, its image headers would have to be written into `SPM.MACS.ABC`, so that other analyses can access these images via `SPM.MACS.ABC.fname` upon loading the `SPM.mat`.[1] Conversely, if you want to implement a new second-level approach using cvLME images from the first level, this method would have to reference `SPM.MACS.cvLME`, so that these images can be accessed via `SPM.MACS.cvLME.fname` upon loading the `SPM.mat`.

On the following pages (see pp. 22-25), we provide code snippets for implementing:

- a method of first-level model assessment taking an SPM-GLM variable `SPM` as an input: In our example, the technique is called ABC, for "A Bayesian Criterion", and we assume that you have written a mathematical function which calculates the ABC over voxels called `function_that_returns_voxel_wise_ABC`.
- a method of second-level model selection taking a model space variable `MS` as an input: In our example, the technique is called DEF, for "Discrete Evidence Fusion", and we assume that you have written a mathematical function which performs DEF in one voxel called `function_that_performs_voxel_wise_DEF`.

Then, the interface and batch functions which have to be added to the MACS toolbox folder would have to look something like this...

---

[1] In MACS toolbox modules belonging to the automatic processing stream, e.g. "MS: perform BMS (automatic)", these images can then be incorporated by simply changing the parameter "Enter LME map" from "cvLME" to "ABC".

#### Snippet 1: interface function for a first-level operation

```
function MA_calculate_ABC(SPM)
% _
% Calculate ABC, A Bayesian Criterion

% Load design
%-----------------------------------------------------------------------%
X = SPM.xX.X;                      % design matrix
K = SPM.xX.K;                      % filtering matrix
W = SPM.xX.W;                      % whitening matrix
V = SPM.xVi.V;                     % non-sphericity

% Load data
%-----------------------------------------------------------------------%
[M m_dim m_ind] = MA_load_mask(SPM);
 Y              = MA_load_data(SPM,m_ind);

% Compute ABC
%-----------------------------------------------------------------------%
ABC         = NaN(size(M));
ABC(m_ind) = function_that_returns_voxel_wise_ABC(Y, X, K, W);

% Save ABC image
%-----------------------------------------------------------------------%
H         =  MA_init_header(SPM, false);
H.fname   = 'MA_ABC.nii';
H.descrip = 'MA_calculate ABC: A Bayesian Criterion';
spm_write_vol(H,reshape(ABC,m_dim));

% Save ABC header
%-----------------------------------------------------------------------%
SPM.MACS.ABC = H;
save(strcat(SPM.swd,'/','SPM.mat'),'SPM');
```

#### Snippet 2: interface function for a second-level operation

```
function MS_perform_DEF(MS)
% _
% Perform DEF, Discrete Evidence Fusion

% Get model parameters
%-----------------------------------------------------------------------%
N = size(MS.SPMs,1);            % number of subjects
M = size(MS.SPMs,2);            % number of models

% Get image dimensions
%-----------------------------------------------------------------------%
load(MS.SPMs{1,1});             % first subject/model
H = spm_vol(SPM.MACS.ABC);      % ABC image header
V = prod(H.dim);                % number of voxels

% Load ABC images
%-----------------------------------------------------------------------%
ABC = zeros(N,M,V);             % N x M x V array of ABCs
for i = 1:N                     % subjects
    for j = 1:M                 % models
        load(MS.SPMs{i,j});
        abc_hdr    = spm_vol(SPM.MACS.ABC);
        abc_img    = spm_read_vols(abc_hdr);
        ABC(i,j,:) = reshape(abc_img,[1 1 V]);
    end;
end;

% Create mask image
%-----------------------------------------------------------------------%
ABC_1 = squeeze(ABC(:,1,:));    % N x V matrix of ABCs for 1st model
[m_img m_hdr m_ind] = MS_create_mask(ABC_1, H);
 v    = numel(m_ind);

% Perform DEF analysis
%-----------------------------------------------------------------------%
DEF = NaN(M,V);
for j = 1:v
    DEF(:,m_ind(j)) = function_that_performs_voxel_wise_DEF(ABC(:,:,j))';
end;

% Save DEF maps
%-----------------------------------------------------------------------%
cd(MS.swd);
for i = 1:M
    H.fname   = strcat(MS.GLMs{i},'_DEF.nii');
    H.descrip = 'MS_perform DEF: Discrete Evidence Fusion';
    spm_write_vol(H,reshape(DEF(i,:),H.dim));
end;
```

#### Snippet 3: batch function for a first-level operation

```
function module = batch_MA_ABC
% _
% Configure MATLAB Batch for MACS Toolbox

% Select SPM.mat
%-----------------------------------------------------------------------%
SPM_mat         = cfg_files;
SPM_mat.tag     = 'SPM_mat';
SPM_mat.name    = 'Select SPM.mat';
SPM_mat.help    = {'Select the SPM.mat file of a specified and/or estimated GLM.'};
SPM_mat.filter  = 'mat';
SPM_mat.ufilter = '^SPM\.mat$';
SPM_mat.num     = [1 1];

% MA: ABC (man)
%-----------------------------------------------------------------------%
module      = cfg_exbranch;
module.tag  = 'MA_ABC';
module.name = 'MA: calculate ABC (manually)';
module.val  = {SPM_mat};
module.help = {'A Bayesian Criterion for General Linear Model'
               'Type "help MA_calculate_ABC" for help.'};
module.prog = @run_module;
module.vout = @vout_module;

% Run batch
%-----------------------------------------------------------------------%
function out = run_module(job)

% execute operation
load(job.SPM_mat{1});
MA_calculate_ABC(SPM);
load(job.SPM_mat{1});
out.ABC_nii = cellstr(strcat(SPM.swd,'/',SPM.MACS.ABC.fname));

% Dependencies
%-----------------------------------------------------------------------%
function dep = vout_module(job)

% define dependencies
dep(1)            = cfg_dep;
dep(1).sname      = 'ABC map (voxel-wise image)';
dep(1).src_output = substruct('.','ABC_nii');
dep(1).tgt_spec   = cfg_findspec({{'filter','image','strtype','e'}});
```

#### Snippet 4: batch function for a second-level operation

```
function module = batch_MS_DEF
% _
% Configure MATLAB Batch for MACS Toolbox

% Select MS.mat
%-----------------------------------------------------------------------%
MS_mat         = cfg_files;
MS_mat.tag     = 'MS_mat';
MS_mat.name    = 'Select MS.mat';
MS_mat.help    = {'Select the MS.mat file describing a model space of GLMs.'};
MS_mat.filter  = 'mat';
MS_mat.ufilter = '^MS\.mat$';
MS_mat.num     = [1 1];

% MS: DEF (auto)
%-----------------------------------------------------------------------%
module      = cfg_exbranch;
module.tag  = 'MS_DEF';
module.name = 'MS: perform DEF (automatic)';
module.val  = {MS_mat};
module.help = {'Discrete Evidence Fusion for General Linear Models'
'Type "help MS_perform_DEF" for help.'};
module.prog = @run_module;
module.vout = @vout_module;

% Run batch
%-----------------------------------------------------------------------%
function out = run_module(job)

% execute operation
load(job.MS_mat{1});
MS_perform_DEF(MS);
load(job.MS_mat{1});
out.DEF_mat = cellstr(strcat(MS.swd,'/','DEF.mat'));

% Dependencies
%-----------------------------------------------------------------------%
function dep = vout_module(job)

% define dependencies
dep(1)            = cfg_dep;
dep(1).sname      = 'DEF results (DEF.mat file)';
dep(1).src_output = substruct('.','DEF_mat');
dep(1).tgt_spec   = cfg_findspec({{'filter','mat','strtype','e'}});
```

**Remark:** In order for these batch modules to appear in the SPM batch editor as part of the MACS toolbox, the batch function names, i.e. `batch_MS_DEF` and `batch_MA_ABC`, have to be added to the variable `module.values` in the function `batch_cfg_master.m`. Before you do this, make a copy of this function under the name `batch_cfg_master_orig.m` in order not to destroy anything.
